# Neuronal migration induces DNA damage in developing brain

**DOI:** 10.1101/2025.06.10.658810

**Authors:** Zhejing Zhang, Andres Canela, Peilin Zou, Takahiro Furuta, Noriko Takeda, Takumi Kawaue, Naotaka Nakazawa, Mai Saeki, Masaki Utsunomiya, Junko Kurisu, Fumiyoshi Ishidate, Hiroyuki Sasanuma, Yusuke Kishi, Mineko Kengaku

## Abstract

Migratory cells tend to have soft nuclei that deform and penetrate narrow spaces^1,2^. Extensive nuclear deformation during migration can cause nuclear envelope rupture and DNA damage in cancer cells, which may contribute to the malignant transformation during tumor progression^3,4,5,6^. However, the significance of DNA damage in physiological migration is less well understood. Here, we demonstrate that the migration of neurons in developing cerebral and cerebellar cortices is accompanied by massive DNA double-strand breaks (DSBs) due to mechanostress during passage through narrow interstitial spaces. Confined migration enhances the binding and cleavage of the genome by topoisomerase IIβ, expressed in neuronal nucleus, independently of the nuclear envelope rupture. Genome sequencing revealed that DSBs tend to occur outside of protein-coding regions and transcription regulatory regions. During normal development, DSBs are rapidly repaired by the non-homologous end joining pathway. The deletion of ligase IV at the onset of neuronal migration leads to persistent DSB accumulation in cerebellar neurons with moderate transcriptional changes in genes related to synaptic function, neuronal development, and stress and immune responses. The mutant mouse develops mild motor deficits in later life, suggesting that the DNA damage generated during normal brain development poses a potential disease risk if left unrepaired.

## Main

Due to their limited regenerative capacity, maintaining genome stability in neurons is crucial for preserving brain function throughout the lifespan^7,8,9,10^. However, neurons are continuously exposed to various sources of DNA damage, including intrinsic factors such as oxidative stress, transcription, and neural activity, as well as extrinsic factors like radiation and environmental toxins^11,12,13^. Neurons are therefore equipped with robust mechanisms to prevent and correct DNA lesions, as excessive or unresolved damages can contribute to brain ageing and neurodegeneration^9,14,15^.

During brain development, newborn neurons migrate from their birthplace in the germinal layers to their final destinations in the emerging cortices and nuclei, where they are integrated into functional neural circuits. Migrating neurons squeeze the nucleus, their largest cargo, through narrow spaces crowded with many other cells and extracellular components^16,17^. Neurons express very low levels of Lamin A, which correlate with the high deformability and migratory capacity of the nucleus^1,18,19,20^. On the other hand, nuclei expressing low Lamin A/C are vulnerable to mechanical stress and risk DNA damage^2,21,22^. In cancer cells and immune cells, severe nuclear deformation during confined migration can cause transient nuclear envelope (NE) rupture and subsequent DNA damage, which has been implicated in cellular senescence and malignant transformation^3,4,5,6,23^. On the other hand, the impact of nuclear deformation during neuronal migration has not been described. In this study, we investigated whether confined migration influences genome integrity of postmitotic neurons during normal brain development.

### Confined migration induces DSBs in granule cells

During postnatal development in mice, cerebellar granule neurons (CGNs) are generated by granule neuron precursors (CGPs) in the outer half of the external granule layer (EGL) (Supplementary Fig. 1a). Postmitotic CGNs move to inner side of the EGL, and then undergo radial migration through the molecular layer (ML) to the internal granule layer (IGL) in the cerebellar cortex during the first three postnatal weeks. The nucleus of migrating neurons exhibits highly dynamic motion, including frequent rotation and deformation^16, 24^. We first observed DNA damage in the developing cerebellar cortex at postnatal day 6 (P6) by immunofluorescence for γH2AX, a marker for DNA double-strand breaks (DSBs), along with the cell proliferation marker Ki67 and the differentiated neuron marker p27Kip1 (Fig. 1a). Many Ki67-positive CGP nuclei in the outer EGL were costained with γH2AX. Some cells with strong γH2AX signals in the ML and IGL were identified as glial cells, as assessed by BLBP expression (Supplementary Fig. 1b).

**Fig. 1:**
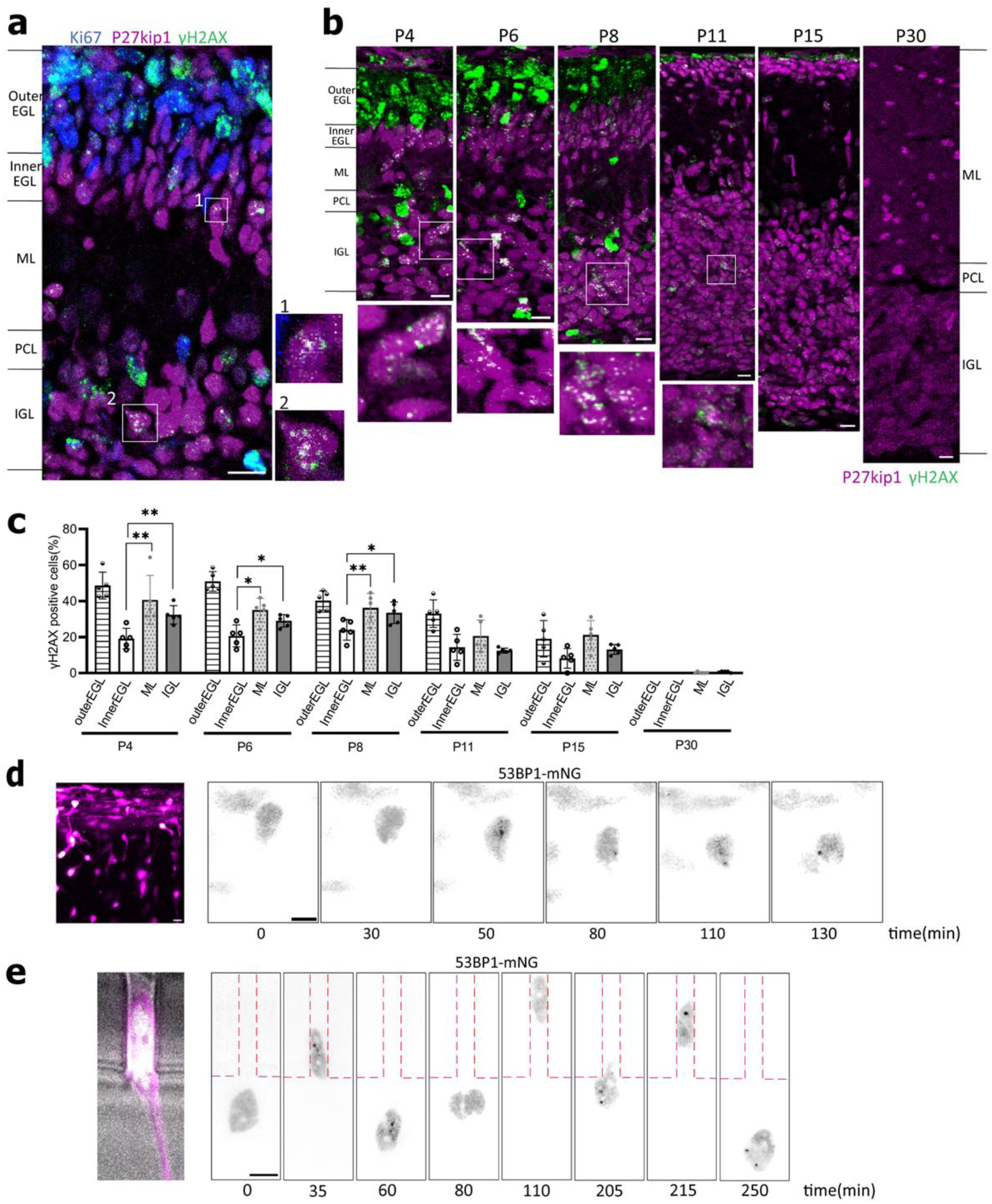
Postmitotic neurons undergo DNA damage by mechanostress during migration through confined interstitial spaces. **a,** Sagittal section of P6 mouse cerebellar cortex stained for γH2AX (green), Ki67 (blue), and p27kip1 (magenta). Postmitotic CGNs in the boxed regions in the ML and IGL that were double stained with p27kip1 and γH2AX are enlarged on the right. **b,** Developing cerebellar cortex from P4 to P30, stained for γH2AX and p27kip1. Boxed regions are enlarged at the bottom. **c,** The percentage of γH2AX-positive cells in the cerebellar cortical layers. Data represent mean ± s.d. *p < 0.05, **p < 0.01, two-way RM ANOVA with Dunnett’s multiple comparisons tests. Samples were collected from 3 independent experiments (n = 5 or 6 mice at each age). **d,** *Left*, Snapshot image of CGNs electroporated with mScarlet (magenta) and 53BP1-mNG (green) in the cerebellar slice from a P7 mouse. *Right*, Image sequence of 53BP1-mNG signals in a CGN migrating from the EGL to the ML in the cerebellar slice culture. **e,** *Left*, A bright-field image of a CGN transfected with mScarlet and 53BP1-mNG. *Right 8 images*, Image sequence of 53BP1-mNG signals during migration in and out of a 3-μm microchannel. Red broken lines outline the channel. Scale bars; **a**, **b**, 10 µm; **d**, **e**, 5 μm.

Besides these DSBs, which were presumably generated by replication stress during cell cycle progression, a significant number of γH2AX foci were observed in postmitotic CGNs in the ML and IGL, costained with p27Kip1 (Fig. 1a).

We then examined γH2AX signals in various developmental stages. Abundant γH2AX signals in the outer EGL were observed at P6 and earlier but sharply declined by P11 as CGPs were exhausted. In contrast, postmitotic CGNs with γH2AX foci in the ML and emerging IGL plateaued from P4 to P8 and persisted until P15 (Fig. 1b). By P30, γH2AX foci were no longer detectable. Considering that it takes ∼72 hours for CGNs to reach the IGL after exiting the cell cycle in the EGL^25^, and that DSB repair occurs in a few hours in normal cells, DSBs in CGNs in the IGL are unlikely the remnant of replication-dependent DSBs generated in CGPs in the outer EGL. Furthermore, the proportion of γH2AX-positive neurons was higher in the ML than in the inner EGL at all examined stages, suggesting that postmitotic CGNs acquire *de novo* DSBs after the onset of radial migration (Fig. 1c). Consistently, γH2AX foci were observed in the nucleus of GFP-positive CGNs migrating in the ML in NeuroD1-GFP transgenic mice (Supplementary Fig. 1c).

To further confirm that DSBs are generated during migration, we electroporated the cerebellum with a DSB marker 53BP1-mNeonGreen (53BP1-mNG) and performed live imaging of CGNs in an organotypic cerebellar slice culture^26^. We observed transient formation of 53BP1-mNG foci in CGNs undergoing radial migration in the ML (Fig. 1d). To ask whether DNA damage is induced by nuclear deformation by mechanical constrictions in dense neural tissue, we monitored dissociated CGNs cultured on microfabricated substrates with constricted channels^27^. CGNs repetitively passed through narrow channels (3 µm in width) while undergoing nuclear deformation and compression. We observed transient accumulation of 53BP1-mNG foci during and after passage of the nucleus through a constriction (Fig. 1e). These results strongly suggest that migrating CGNs undergo DSBs due to mechanostress while migrating in a confined environment in the developing brain.

We next examined whether DSB formation during neuronal migration is fundamental among neuronal types, using cerebellar Purkinje cells and neocortical neurons. Differentiated Purkinje cells migrate from the ventricular zone (VZ) to the cerebellar primordium at embryonic day 12-18 (Supplementary Fig. 1d)^28^. γH2AX foci were detected in the nuclei of postmitotic Purkinje cells labeled by anti-Lhx1/5 during and after migration. Additionally, we observed significant γH2AX foci in emerging neocortex at E15. The number of γH2AX-positive cells was high in the VZ populated by neural progenitor cells and declined in the intermediate zone (IZ) as differentiation proceeds.

Nuclei with γH2AX foci increased again in the subplate (SP), suggesting *de novo* DSB formation in postmitotic neurons after the onset of radial migration (Supplementary Fig. 1e). Live-imaging of cortical neurons in an organotypic culture also revealed the transient formation of 53BP1-mNG foci during radial migration in the cortical plate (Supplementary Fig. 1f). Together, these observations indicate that at least three types of postmitotic neurons undergo DNA damage during neuronal migration in the developing brain.

### DSB during migration involves TOP2-dependent cleavage independent of NE rupture

In cancer cell lines, DNA damage during confined migration has been shown to be exacerbated by NE rupture, which permits the entry of cytoplasmic nuclease and the leakage of DNA damage repair factors from the nucleus^5,6^. We confirmed NE rupture events in HeLa cells using a fluorescent reporter fused to a nuclear localization sequence (mNG-NLS) that rapidly spread in the cytoplasm as cells migrated through constricted channels^3,4^(Fig. 2a). In contrast, in CGNs cultured in the same patterned dish, NE rupture was never observed, even in severely deformed nuclei migrating through narrow spaces (Fig. 2b). HeLa cells expressing catalytically inactive cGAS fused to GFP (GFP-cGAS), a marker of NE rupture^4,21,29^, showed GFP-cGAS foci in the nuclear periphery after transmigration through 3-μm pores in a transwell. In contrast, peripheral accumulation of GFP-cGAS was not observed in CGN nuclei with or without confined migration in transwells (Fig. 2c). NE blebbing and rupture were rare events seen in only a few percent of CGNs migrating through 3-μm pores, as indicated by immunofluorescence with a nuclear lamina marker Lamin B1 and a chromatin marker H4K20me3 (Supplementary Fig. 2a, b). These findings suggest that, unlike cancer cells, CGNs generate DNA damage by mechanical stress during confined migration independently of NE rupture.

**Fig. 2:**
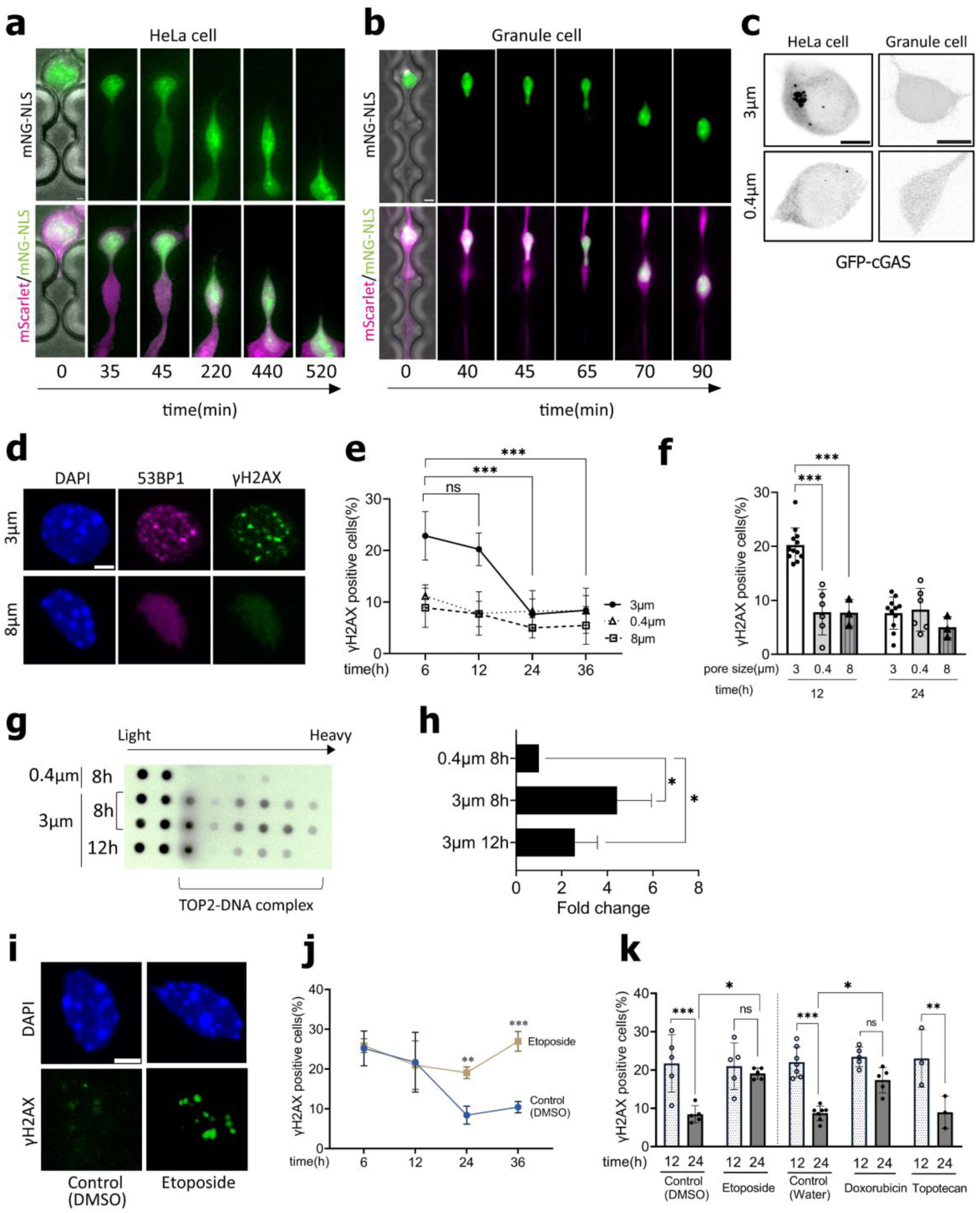
DNA damage during neuronal migration is associated with Top2-dependent cleavage. **a**, **b**, A bright field image (left) and timelapse sequences (right 5 images) of a HeLa cell (**a**) and a CGN (**b**) transfected with mScarlet and mNG-NLS migrating through 3-μm microchannels. **c**, HeLa cells (left) and CGNs (right) transfected with cGAS-GFP. Upper images are the cells after transwell migration through 3-μm pores and lower images are non-migrated cells on 0.4-μm pores. **d**, Immunostaining of CGNs migrated through 3-μm or 8-μm pores at 12 h after seeding. **e**,**f**, The percentages of γH2AX-positive nuclei after transwell migration through confined (3-μm) and non-confined (8-μm) pores, and non-migrated nuclei on 0.4 µm pores, at indicated time after inoculation. **g**, Measurement of TOP2β-DNA cleavage complex formation during migration. CGNs migrated through 3-μm pores and non-migrated cells on 0.4-μm pores were collected at 8 or 12 hours after seeding. Genomic DNA isolated from CGNs in transwell cultures was subjected to sedimentation by the CsCl-gradient ultracentrifugation. Individual fractions were blotted and subjected to immunoblot analysis with TOP2β antibody. The lightest two fractions include free TOP2, while the other heavier fractions include TOP2cc. **h**, Quantification of TOP2cc in the CGNs at 8 h and 12 h after 3-µm pore migration relative to the amount of TOP2cc in non-migrated CGNs on 0.4-µm pores. **i**, CGNs migrated through 3-μm pores in the presence or absence of etoposide. Cells were fixed at 24 h after seeding and stained with γH2AX. **j**, Inhibition of time-dependent decay of the γH2AX-positive nuclei after 3-µm pore migration by the treatment with etoposide. **k**, The effect of topoisomerase inhibitors on the percentage of γH2AX-positive nuclei at 12 h and 24 h of transwell assay through 3-µm pores. **e, f, h, j, k,** Samples were collected from at least 3 independent experiments per group. **d, e, j, k,** two-way ANOVA with Tukey’s multiple comparisons tests; **h,** two-tailed unpaired t-tests. Data represent mean ± s.d. *p < 0.05, **p < 0.01, ***p < 0.001. Scale bars; **a-c**, 5 μm; **d**, **i**, 3 μm.

To investigate the mechanism underlying transient DNA damage in CGNs during confined migration, we adopted a transwell assay with different pore sizes^27^. The majority of the CGNs reached the bottom surface within 12 hours (h) of seeding through both confined 3-μm pores (56%) and permissive 8-μm pores (79%) (Supplementary Fig. 2c). We observed a significantly higher proportion of cells generating γH2AX foci after confined migration through 3-μm pores, compared to those after non-confined migration through 8-μm pores or unmigrated cells on top of surfaces with 0.4-μm pores (Fig. 2d-f). Similar results were obtained for 53BP1 staining (Fig. 2f, Supplementary Fig. 2d, e).

Consistent with previous reports using super-resolution imaging, we confirmed that γH2AX foci in postmigratory CGNs form characteristic nanodomains in both cerebellar cortex and transwell cultures, typically clustering around a single spot of Ku70, which is involved in DSB repair via the nonhomologous end joining (NHEJ) pathway (Supplementary Fig. 2f) ^30,31^. More than 20% of CGNs bore multiple γH2AX and/or 53BP1 foci at 6 h and 12 h after transmigration through 3-μm pores, but the levels decreased to the control level at 24 h and thereafter (Fig. 2d, e; Supplementary Fig. 2d, e). TUNEL assay revealed little to no increase in apoptotic cell death up to 36 h, suggesting that DSBs induced by confined migration are repaired without causing apoptosis (Supplementary Fig. 2g).

Since neuronal DSBs induced by confined migration appeared to occur independently of NE rupture, they are thought to result from mechanical strain on the chromatin structure. We predicted the involvement of Topoisomerase II (TOP2), a key enzyme that relieves torsional stress during replication and transcription. TOP2 catalyzes a temporary DSB in a DNA molecule to allow the passage of another DNA segment, followed by religation. During this process, TOP2 becomes covalently attached to the ends of the DNA break, forming a transient intermediate called TOP2-DNA cleavage complex (TOP2cc). Failure of TOP2 to religate the break and complete its catalytic cycle results in the accumulation of persistent TOP2ccs, which must then be resolved through proteasomal degradation and DNA repair^32^. In agreement with previous reports, TOP2β was highly expressed in the nucleus of postmitotic CGNs undergoing radial migration (Supplementary Fig. 2h)^33^. We measured the transient TOP2βcc in CGNs in a transwell assay^34^. In unmigrated cells on 0.4 µm pore filters, free TOP2β unbound to DNA and a very small amount of TOP2βcc were detected. In contrast, strong Top2βcc signals were detected in CGNs that migrated through 3-μm pores at 8 h after seeding (Fig. 2g). TOP2βccs were rapidly decreased by 12 h, suggesting that TOP2β fails to complete DNA religation, becomes trapped on DNA, and is subsequently repaired during and after confined migration (Fig. 2h). We next examined the effect of TOP2 inhibitors etoposide and doxorubicin, which strongly trap TOP2 in the TOP2cc state by preventing DNA ligation after cleavage. Neither drug altered DSB formation in transmigrated CGNs at 6 h and 12 h. However, DSBs were persisted until 36 h at levels comparable to those observed at 12 h, supporting that the DSBs formed during migration are induced by TOP2 (Fig. 2i-k; Supplementary Fig. 2i). In contrast, Topotecan, a TOP1 poison stabilizing TOP1cc, had minimal effect on γH2AX reduction at 24 h (Fig. 2k; Supplementary Fig. 2i). These findings suggest that nuclear deformation during confined migration causes structural change in chromatin, leading to TOP2β trapping and subsequent cleavage of DNA strands in neurons.

### DSB during confined migration is repaired via the NHEJ pathway

DSBs in CGNs after transmigration through constricted 3-μm pores were mostly depleted by 24 h without causing cell death (Fig. 2d; Supplementary Fig. 2g). Consistently, the abundant DSBs in CGNs also disappeared without apparent signs of apoptosis in the cerebellum, suggesting that CGNs successfully repair these DSBs during development (Supplementary Fig. 3a). Previous studies have indicated that TOP2-dependent DSBs at sites of activity-dependent genes are repaired by the NHEJ pathway in CNS neurons^35^.

We thus asked if the NHEJ pathway is involved in migration-induced DSB repair. In a transwell assay, treatment with SCR7 or NU7441, inhibitors of the key NHEJ components Ligase IV (Lig4) and DNA-PK respectively, disrupted the decay of γH2AX foci between 12 and 24 h after transmigration through 3-μm pores. In contrast, inhibition of the homologous recombination pathway by a Rad51 inhibitor B02 had minimal impact on γH2AX foci reduction (Fig. 3a, b).

**Fig. 3:**
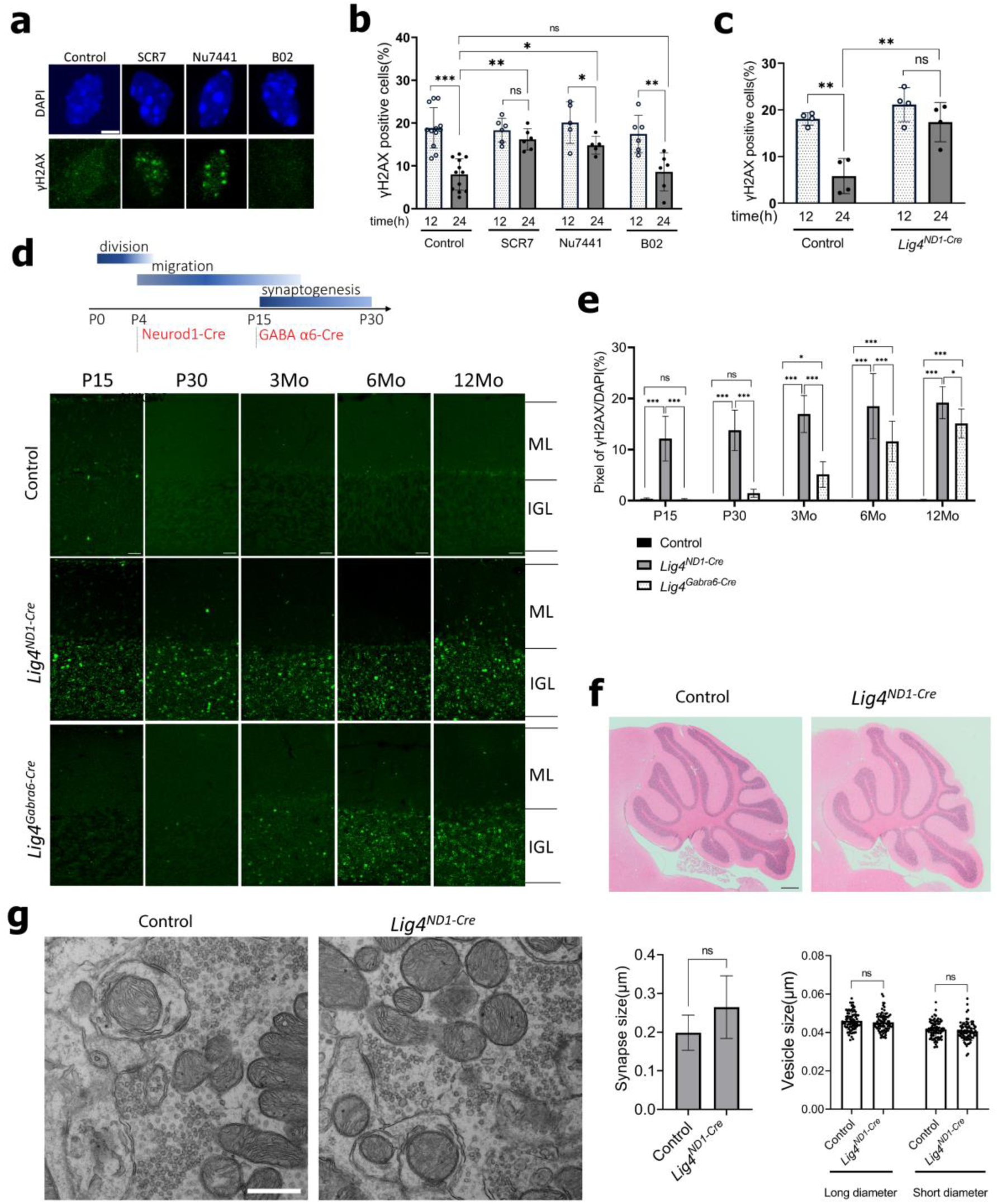
DNA damage during neuronal migration is repaired by the NHEJ pathway. **a**, The effects of DSB repair inhibitors on migration-induced γH2AX foci. Cells were fixed at 24 h after seeding on 3-μm pore filters. **b**, The percentage of γH2AX-positive CGNs treated with DSB repair inhibitors at 12 h and 24 h in transwell culture through 3-µm pores. Samples were collected from more than 3 independent experiments per group. **c**, γH2AX-positive nuclei from control (Neurod1-Cre^-/-^; Lig4^flox/flox^) and *Lig4 ^ND^*^1^*^-Cre^*mice in 3-µm transwell assays. Samples were collected from 4 independent experiments. **d**, Timing of Neurod1 and GABAα6 expression in developing CGNs (top) and γH2AX foci formation in the cerebellar cortices from control, *Lig4 ^ND^*^1^*^-Cre^*and *Lig4 ^Gabra^*^6^*^-Cre^* mice (bottom). **e**, Quantification of γH2AX foci (relative pixel intensities of γH2AX and DAPI signals) in the IGL. n = 5 mice per group. **f**, Hematoxylin and eosin staining of the cerebellar cortex from control and *Lig4 ^ND^*^1^*^-Cre^*mice at 12 mo. **g**, Electron microscopy images (right) and quantification of the synapse and vesicle sizes in the IGL of control and *Lig4 ^ND^*^1^*^-Cre^*cerebella at 14 mo. n = 3 mice per group. **b**, **c**, **e**, two-way ANOVA with Tukey’s multiple comparisons tests. **g,** two-tailed unpaired t-tests. Data represent mean ± s.d. *p < 0.05, **p < 0.01, ***p < 0.001, Scale bars; **a**, 3 μm; **d**, 24 μm. **f**, 500 μm**. g**, 500 nm

To further investigate the role of the NHEJ pathway, we analyzed the effect of conditional deletion of Lig4 in postmitotic CGNs (Neurod1-Cre^+/-^; Lig4^flox/flox^, hereafter called *Lig4^ND^*^1^*^-Cre^*)^36^. CGNs from *Lig4^ND^*^1^*^-Cre^*mice exhibited impaired DSB repair in transwell assays, with persistent γH2AX foci at 24 h (Fig. 3c). In the cerebellar cortex of *Lig4 ^ND^*^1^*^-Cre^* mice, abundant γH2AX foci were detected in the IGL at P15 and P30, whereas in control mice (Neurod1-Cre^-/-^; Lig4^flox/flox^), the foci had largely disappeared by these stages (Fig. 1b and 3d). γH2AX accumulation in the cerebellum persisted until 12 months (mo) in *Lig4 ^ND^*^1^*^-Cre^* mice. To verify that these γH2AX foci represent unrepaired DSBs generated during migration, we made another Lig4 deficient mice using a GABAα6-Cre driver (*Lig4 ^Gabra^*^6^*^-Cre^*), in which Lig4 is depleted in postmigratory CGNs in the IGL^37^.

Although Lig4 deletion is expected to occur in postmigratory CGNs as early as P10, γH2AX accumulation was not seen until 3 mo in *Lig4 ^Gabra^*^6^*^-Cre^*mice (Fig. 3d). γH2AX levels gradually increased with age, likely due to slowly accumulating unrepaired DSBs generated by neuronal activity and other stressors. However, the levels of γH2AX accumulation remained significantly higher throughout life in *Lig4 ^ND^*^1^*^-Cre^* mice, which drive Lig4 deletion in CGNs before the onset of migration only one week earlier than *Lig4 ^Gabra^*^6^*^-Cre^*mice (Fig. 3e). These results suggest that mechanostress during confined migration is a primary trigger of DNA damage in juvenile CGNs during development, rather than other stressors like transcription or neural activity. It is also indicated that the DNA damage during CGN migration is rapidly repaired via the NHEJ pathway.

Unexpectedly, the persistent DNA damage caused no obvious morphological abnormalities or cell death in the cerebellum of *Lig4 ^ND^*^1^*^-Cre^* mice, as observed by both light and electron microscopies (Fig. 3f, g; Supplementary Fig. 3b, c). Thus, CGNs can tolerate DSBs generated during and after migration without major impacts on cellular functions.

### DSBs are broadly distributed outside regions of active chromatin

Next, we mapped the positions of DSBs induced by confined migration using END-seq, a sequencing-based method that monitors DSBs genome-wide at base-pair resolution^38^.

CGNs were subjected to transwell migration through 3-µm pores in the presence of etoposide and were collected at 6 h from the bottom. CGNs after confined migration exhibited more peaks than unmigrated control (Fig. 4a). Upon etoposide treatment, which traps TOP2 at the cleavage stage, the END-seq signal serves as a readout of TOP2 activity in the genome. As previously reported, we observed an enrichment of END-seq signals at sites co-occupied by both CTCF and RAD21 in both migrated and unmigrated cells (Supplementary Fig. 4a,b)^39,40^. However, aside from CTCF/RAD21-bound sites, no consistent etoposide END-seq breaks were detected at specific motifs or gene loci in transmigrated cells. Given this lack of site-specific DSBs, we instead examined broader genomic preferences by classifying END-seq peaks according to seven chromatin categories: active promoters, active enhancers, active exons, active introns, transcriptionally inactive and unlabeled euchromatin, lamina-associated domains (LADs), and heterochromatin not overlapping LADs. These regions were defined based on chromosome accessibility (ATAC-seq), histone modifications (H3K4me3 and H3K9me3 ChIP-seq), nuclear lamina association (Lamin B1 ChIP-seq), and transcriptional activity (RNA-seq), from unmigrated control CGNs. We then profiled the landscape of DSBs that occurred in CGNs with or without confined migration. Consistent with the association of TOP2 cleavage with active chromatin, END-seq signals common to both migrated and unmigrated CGNs were enriched in open chromatin, including active promoters and enhancers (Fig. 4b). In contrast, DSBs generated specifically by confined migration were largely excluded from the active transcription regulatory elements and instead occurred more frequently in introns, transcriptionally inactive euchromatin, and LADs (Fig. 4b). Migration-induced DSBs tended to overlap more frequently with repetitive DNA particularly retrotransposons LINEs and LTRs (Fig. 4c). To examine the relationship between etoposide-induced DSBs and transcriptional activity, we analyzed END-seq signal over all transcription start sites (TSS) and gene bodies. Confined migration caused little or no change in DSB levels at promoters or within gene bodies, regardless of transcriptional activities based on RNA-seq (Supplementary. 4c). Similarly, no significant difference was observed in long genes either, which have been associated with neuronal physiology and connectivity^41^ (Supplementary Fig. 4d). We also performed END-seq using cerebellar tissues from P15 control and *Lig4 ^ND^*^1^*^-Cre^* mice to map the unrepaired DSBs in the NHEJ pathway-deficient condition in vivo. As in the transwell system, the DSBs in *Lig4 ^ND^*^1^*^-Cre^* mice also lacked site-specificity and were enriched in chromatin categories classified as inactive chromatin and retrotransposons (Supplementary Fig. 5a-c). No accumulation was observed in the promoters or gene bodies of transcribed genes (Supplmentary Fig.5d and e). These results suggest that DSBs generated during confined migration are topologically and functionally different from the typical TOP2β hotspots, which include regulatory elements and chromatin loop anchors.

**Fig. 4:**
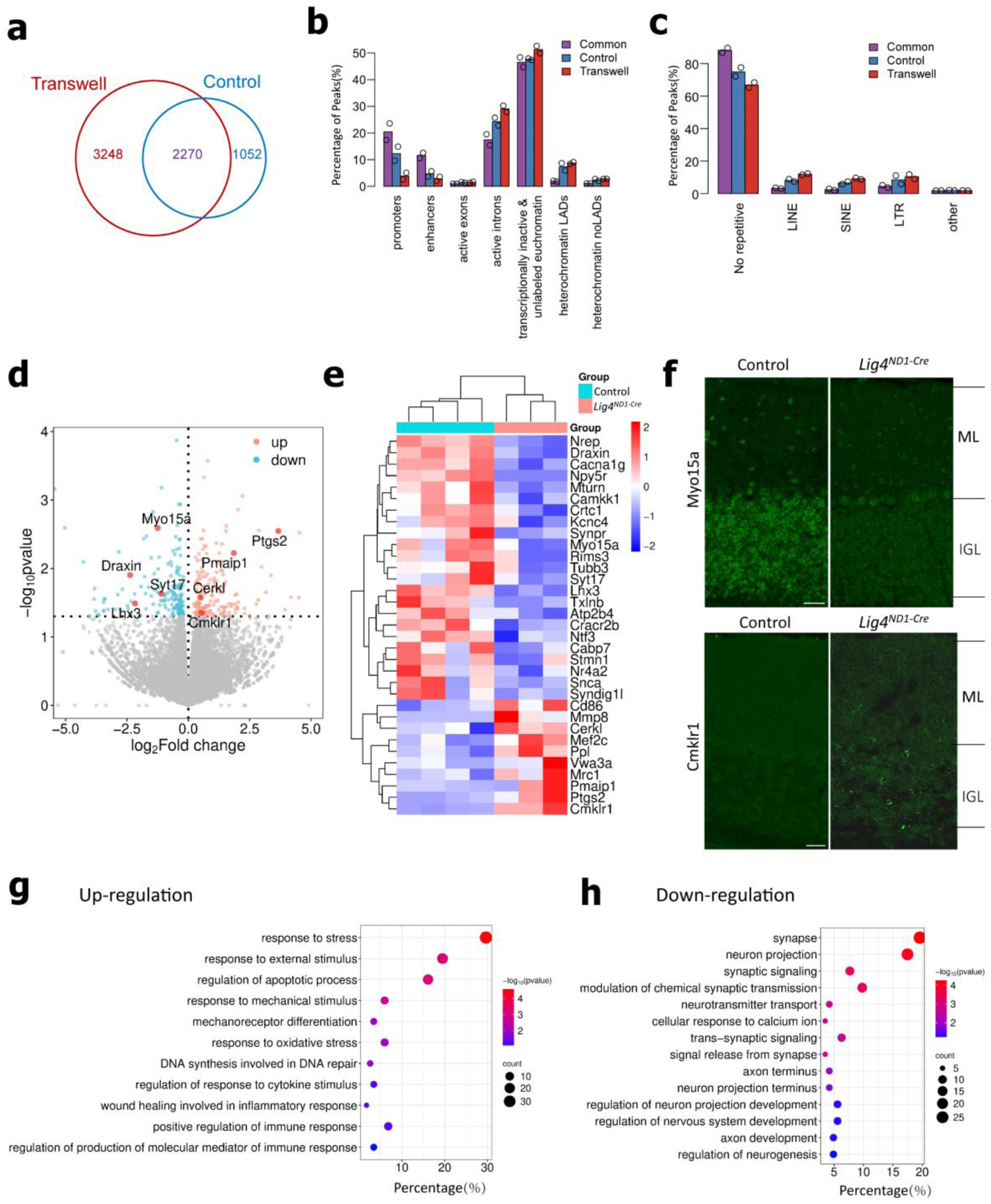
Persistent DSBs in postmigratory neurons lead to transcriptional changes. **a**, Venn diagram showing the overlap of END-seq peaks (etoposide-induced TOP2 lesions) in CGNs after transwell migration through 3-µm pores and unmigrated CGNs on culture dishes. **b,** Barplot comparing the relative frequency of specific END-seq peaks across defined chromatin categories (see Methods); and **c**, Barplot comparing the relative levels of specific END-seq peaks in repetitive elements. Bars represent average and circles indicate individual biological replicas (n = 2). **d**, Transcriptional changes in cerebellar tissue between 2 mo control and *Lig4 ^ND^*^1^*^-Cre^*mice (n = 4 mice for control, n=3 mice for *Lig4 ^ND^*^1^*^-Cre^*). Colored dots indicate significant differentially expressed genes (DEGs). **e**, Heatmap of DEGs related to neuronal function and development, stress and immune response, color refers to different Z-scores calculated from individual RPKM. **f**, Immunostaining of Myo15a (top) and Cmklr1(bottom) in the cerebellar cortex of control and *Lig4 ^ND^*^1^*^-Cre^* mice at 3 mo. Scale bar, 24 μm. **g**, **h**, Upregulated (**g**) and downregulated (**h**) pathways detected via Gene Ontology (GO) enrichment analysis using DEGs. The horizontal axis represents the percentage (%), and the vertical axis lists the enriched GO category names. The color scale represents different -log10 (p-value) thresholds, and dot size corresponds to the number of genes associated with each pathway.

In an attempt to clarify the long-term influence of unrepaired DSBs in Lig4-deficient cerebellum, we performed unpaired bulk RNA-seq on cerebellar lysates from control and *Lig4 ^ND^*^1^*^-Cre^* mice at 2 mo. Although overall differences between control and *Lig4 ^ND^*^1^*^-Cre^*were not huge, we identified 336 differentially expressed genes (DEGs) among 50,531 genes (Fig. 4d). Genes related to neuronal differentiation, including Draxin, LIM homeobox 3 (Lhx3), myosin 15a (Myo15a) and synaptotagmin 7 (Syt7), showed significant downregulation, while those related to stress response and immune response which primarily expressed in immune cells and microglia, such as prostaglandin-endoperoxide synthase 2 (Ptgs2), phorbol-12-myristate-13-acetate-induced protein 1(Pmaip1) and chemerin chemokine like receptor 1 (Cmklr1), were upregulated (Fig. 4e). For validation, we performed immunofluorescence of the genes showed expression changes in the analysis and confirmed the downregulation (Myo15a) and upregulation (Cmlkr1) in the IGL of *Lig4^ND^*^1^*^-Cre^* cerebellum (Fig. 4f). Gene ontology (GO) analysis further confirmed the downregulation of genes involved in synapse formation and function, as well as the upregulation of stress- and immune-related genes in *Lig4 ^ND^*^1^*^-Cre^* mice (Fig. 4g, h). Comparison with public datasets showed that these gene expression changes were not typical of normal cerebellar ageing (Supplementary Fig. 6a-c)^42,43,44^.

However, the upregulated gene cluster in Lig4 *^ND^*^1^*^-Cre^*overlapped with those upregulated in the cerebrum in the early phase of the CKp25 mouse, a model of neurodegeneration, suggesting a link between DNA damage and a senescence-like state in neurodegenerative diseases (Supplementary Fig. 6d)^45^.

### Persistent DSBs by Lig4-deficiency leads to mild cerebellar dysfunction

Although *Lig4 ^ND^*^1^*^-Cre^* mice appeared to develop normally, they gradually exhibited mild motor discoordination. With age, adult *Lig4 ^ND^*^1^*^-Cre^*mice were observed to crawl from 12 mo and onward (Fig. 5a). In a footprint test, forelimb-hindlimb coordination was disrupted in *Lig4 ^ND^*^1^*^-Cre^*mice, such that the average width between the right and left hindlimb gaits was increased, whereas that of forelimbs remained comparable to wildtype littermates (Fig. 5b). Additionally, the stride length of hindlimbs became shorter and more irregular in *Lig4 ^ND^*^1^*^-Cre^*mice (Fig. 5b, c). We also conducted a balance beam test across different ages. *Lig4 ^ND^*^1^*^-Cre^*mice exhibited increased hindlimb slips when crossing a narrow beam from 3 mo onward, which worsened over time. In contrast, *Lig4 ^Gabra^*^6^*^-Cre^* mice performed similarly to control at 3 mo but progressively developed deficits with age (Fig. 5d). *Lig4^ND^*^1^*^-Cre^* mice displayed a limited number of γH2AX foci in multiple areas which may affect to motor coordination, including the cortex, thalamus, and subthalamic nucleus (STN) from 3 mo (Supplementary Fig. 7a, b). However, *Lig4^Gabra^*^6^*^-Cre^*mice gradually developed motor deficits without forming γH2AX accumulation outside the cerebellar system. The cerebellar IGL was the only area in the brain where DSB accumulation was high in both *Lig4 ^ND^*^1^*^-Cre^* and *Lig4 ^Gabra^*^6^*^-Cre^*, suggesting that the altered behaviors are caused by CGN dysfunction (Supplementary Fig. 7a, b). Performance on a rotating rod was comparable between control and Lig4-deficient mice, suggesting motor learning is not affected by sustained DSBs (Fig. 5e). Body weight and grip strength remained unaffected in all groups, confirming that balance beam deficits were not due to muscle weakness (Supplementary Fig. 7c, d). These findings suggest that persistent DNA damage in Lig4-deficient CGNs leads to mild but progressive motor discoordination.

**Fig. 5:**
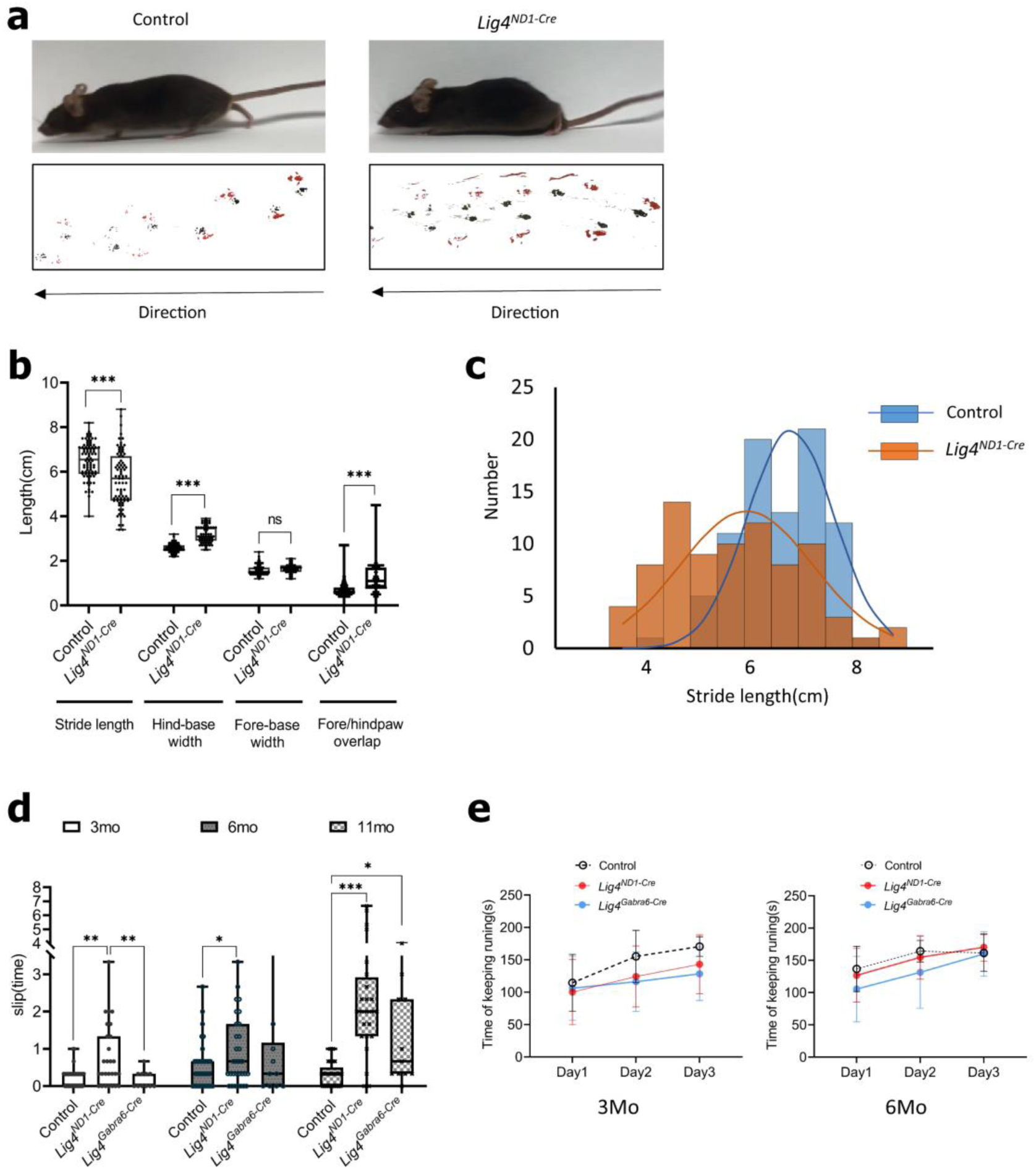
Progressive motor deficits in mice with persistent DSBs in CGNs. **a**, Representative walking posture (top) and footprints (bottom) of control (left) and *Lig4 ^ND^*^1^*^-Cre^*(right) mice at 12 mo. Gait patterns of fore paws (black) and hind paws (red) shown. **b**, Quantitative comparison of walking behavior. *Lig4 ^ND^*^1^*^-Cre^*mice showed a greater hind-base width, shorter stride length, and greater fore and hind paw distance. n > 10 mice per group. **c**, Histogram of stride length distribution of control and *Lig4 ^ND^*^1^*^-Cre^* mice. **d**, Balance on the raised round beams. The number of hindlimb slips was recorded on three independent trials. Mice of indicated genotypes were tested at three different ages. n ≥ 10 mice per group. **e,** Quantification of time spent running on the rotarod over three consecutive days at 3 and 6 mo (n ≥ 7 mice for control; n > 10 for *Lig4 ^ND^*^1^*^-Cre^*; n = 4–6 for *Lig4 ^Gabra^*^6^*^-Cre^*). Data represent mean ± s.d. *p < 0.05, **p < 0.01, ***p < 0.001, Welch’s t-tests.

## Discussion

In this study, we demonstrate that newly born neurons inevitably undergo DNA damage by mechanostress during migration. The transient DSBs are faithfully repaired during normal brain development without causing obvious defects in the mature brain. Unlike in multiple cancer cell lines, which undergo NE rupture and allow nuclear entry of cytoplasmic nucleases^5,6^, cerebellar neurons rarely experience NE rupture and instead form DSBs through TOP2 activity. Thus, DSB formation in migrating neurons appears to be more systematic than those induced by unexpected entry of cytoplasmic nucleases.

TOP2 relaxes torsional stress in the DNA by catalyzing transient DSBs. Extensive studies have demonstrated the binding of TOP2 to intertwined DNA in front of the replication fork^46^ or of the transcription machineries^47^, and at the CTCF/Cohesin loop anchors^39^.

Contrary to our initial prediction that the mechanostess during migration may enhance TOP2-dependent cleavage at loop anchors or transcription initiation sites, the damage appeared independent of these TOP2 target sites. Instead, they tended to be found in ‘quieter’ and ‘safer’ places, including protein non-coding regions and transcriptionally inactive chromatin. Interestingly, TOP2α breaks during replication have been shown to enrich in heterochromatin and repetitive sequences including LTRs and LINEs, supporting the idea that these regions could be native TOP2 targets for decatenation or resolving topological stress^48^. Alternatively, it may be that migration-induced TOP2 cleavages are generated in both active and inactive chromatin; however, processing of TOP2cc into DSBs is faster in transcribed regions^40^, making those in active chromatin more rapidly resolved than those in inactive chromatin. In addition, DSBs in inactive chromatin are repaired more slowly due to the lower efficiency of DNA repair compared to actively transcribed euchromatin^49^. The regulatory mechanism of TOP2 targeting is yet to be clarified, but topological changes in DNA strands by bending or torsion are thought to affect the steric accessibility and activity of TOP2^50,51^. Stretching and compression of chromosomes by large nuclear deformation during confined migration may lead to strain accumulation in multiple genome regions, where TOP2 binds and alleviates excessive tensile and torsional stress on DNA during confined migration. It is also known that mechanical distortion of chromatin could impair TOP2’s ability to religate DNA strands after cleavage^52^, leading to increased formation of TOP2 cleavage complexes and DNA damage.

Transcriptomic analysis of *Lig4^ND^*^1^*^-Cre^* mice revealed only modest changes with slight downregulation of genes related to neuronal development and function, and an upregulation of stress and immune response genes. Given that many upregulated genes are predominantly expressed in non-neuronal cells and that the DSB sites are not closely associated with these genes, the transcriptional changes are unlikely to be a direct consequence of structural alterations in transcription regulatory regions by DNA breaks. It has been suggested that persistent DSBs induce pro-inflammatory signals in neurons, which in turn activate immune response genes in nearby microglia in CKp25 mouse model of neurodegeneration^53^. Although the transcription changes are much smaller and the upregulation of pro-inflammatory signals in neurons, such as NFκB, was not evident in our *Lig4^ND^*^1^*^-Cre^*mice, there was significant overlap in the upregulated gene clusters between CKp25 mice and *Lig4^ND^*^1^*^-Cre^*mice. We surmise that persistent DSBs in neurons lead to mild inflammatory conditions in brain tissue, which in turn cause the downregulation of neuronal genes in *Lig4^ND^*^1^*^-Cre^*mice. The progressive mild motor discoordination without apparent cell death or anatomical defects seen in *Lig4^ND^*^1^*^-Cre^* mice might be a consequence of these modest gene expression changes. Our results suggest that unlike mitotic DNA damage, which can severely alter transcription, differentiation and even lead to cell death if left unrepaired^54,55,56^, migration-induced DNA damage in postmitotic CGNs has limited impacts on genome stability and function. This aligns with the notion that the damage is confined to safe regions of the genome, sparing critical gene loci and minimizing the overall impact on neuronal function.

We identified the NHEJ pathway as the primary repair mechanism for migration-induced DSBs. The NHEJ repair involves the direct ligation of DNA ends, potentially causing small insertion and deletions^57,58^. Robust DSBs during migration may thus cause somatic mosaicism, which is abundant in the brain and may link to neuronal individuality as well as neurodevelopmental and neurodegenerative diseases^8,10,13,59,60,61^. It is also noteworthy that progressive cerebellar motor deficits are a common symptom in patients with genome instability syndromes caused by defective DNA repair pathways^14,15,62^. The vulnerability of neurons to DNA repair defects has been attributed to frequent DNA damage caused by high oxidative metabolism and transcriptional activity. It is, however, not clear why the cerebellum is particularly susceptible to late-onset impairments.

Whether the DSB during neuronal migration are related to the pathogenesis of genome instability syndromes remains unclear, given the marked pathological differences, such as the presence or absence of severe neurodegeneration. Nonetheless, our data suggest that mechanostress during neuronal migration represents one of the major sources of endogenous DNA lesions in the cerebellum, which may be involved in pathogenesis of diseases characterized by cerebellar dysfunction.

## Methods

### Materials

#### Antibodies

The antibodies used are as follows: 53BP1(Novous, NB100-304, 1:1000 for immunofluorescence); γH2AX(Merck, 05-636, 1:1000 for immunofluorescence): γH2AX (CST, 9718S, 1:300 for immunohistochemistry); P27Kip1(BD Transduction Laboratories, 610241, 1:400 for immunohistochemistry and immunofluorescence); Ki67(eBioscience, 14-5698-80, 1:500 for immunohistochemistry or 1:1000 immunofluorescence); TOP2β (Novous, NB100-40842, 1:500 for immunohistochemistry or 1:10000 For Western blot); Ku70 (Abcam, ab61783, 1:400 for immunofluorescence); BLBP (Cosmo Bio, MSFR100310, 1:100 for immunohistochemistry); Lhx1/5 (Hybridoma Bank, 1:5 for immunohistochemistry); Cleaved-caspase3 (CST, 9579S, 1:100 for immunohistochemistry); Vglut1(Merck, AB5905, 1:1000 for immunohistochemistry); Vglut2(Merck, AB2251-I, 1:1000 for immunohistochemistry); H4K20me3(Abcam, ab78517, 1:500 for immunofluorescence); H3K4me1 (Abcam, ab8895, 10ug for ChIP); H3k27ac (Abcam, ab4729, 10ug for ChIP); H3K4me3 (Merck, 07-473, 10ug for ChIP); H3K9me3 (Abcam, ab8898, 10ug for ChIP); CTCF (Merck 07-729, 10ug for ChIP); Rad21(Abcam, 05-908, 10ug for ChIP); LaminB1 (Abcam, ab16048, 1:500 immunofluorescence or 10ug for ChIP); Myo15a (Novous, NOV-NBP2-69069, 1:100 for immunohistochemistry); Cmklr1(Bioss Inc, BIS-BS-10185R-TR, 1:200 for immunohistochemistry); Alexa Fluor 488, 555, 647 secondary antibodies (Thermo Fisher Scientific, 1:1000)

#### Chemicals

Etoposide (Sigma-Aldrich E1383, 1nM); Doxorubicin (Cayman Chemical 15007, 1nM); Topotecan (Cayman Chemical 14129, 300nM); SCR7 (Cosmo Bio M60082-2S, 100 μM); NU7441 (SelleckChem S2638, 20μM); B02 (Cayman Chemical CAY-22133, 1.2μM); Palbociclib (Sigma-Aldrich PZ0383, 5μM)

#### Plasmids

The pCAG-mScarlet plasmid was generated using the mScarlet sequence obtained from Addgene (#85042; a gift from Dorus Gadella, University of Amsterdam). The pCAG-mNeonGreen-NLS plasmid was constructed by inserting the mNeonGreen sequence from pmNeonGreen-N1, fused with a nuclear localization signal (NLS), into the pCAGGS backbone, as previously described^27^. To generate the pCAG-53BP1-mNeonGreen construct, mouse 53BP1 cDNA was amplified from a mouse brain cDNA library and inserted into the pENTR1A vector. In parallel, the Gateway cassette from pDest -eGFP-N1 was inserted into the EcoRI site of pmNeonGreen using In-Fusion Cloning to generate pDest-mNeonGreen. The Gateway cassette containing mNeonGreen was then inserted into the EcoRI site of pCAGGS to generate pCAG-Dest-mNeonGreen, also via In-Fusion Cloning. Finally, 53BP1 cDNA in pENTR1A was recombined into pCAG-Dest-mNeonGreen using LR Clonase II Enzyme Mix (Invitrogen, 11791020) to produce the final pCAG-53BP1-mNeonGreen construct. The pTRIP-CMV-GFP-FLAG-cGAS plasmid was obtained from Addgene (#86674; a gift from Nicolas Manel). The pAAV-Neurod1-GFP plasmid was constructed by replacing the CAG promoter in pAAV-CAG-EGFP^63^ with the Neurod1 promoter, which was cloned from the cDNA of CGNs isolated from P6 ICR mice.

#### Animals

All animal experiments were approved by and conducted in accordance with the guidelines of the Animal Experiment Committee of Kyoto University. For some in vitro experiments, timed-pregnant C57BL/6J and ICR mice were purchased from Japan SLC. We generated conditional deletion mice by breeding *Lig4 ^flox/flox^* ^36^ with *NeuroD1-Cre* (STOCK Tg (Neurod1-cre) RZ24Gsat/Mmcd, MMRRC_036320-UCD) or *Gabra6-Cre* (B6.129P2-Gabra6^tm2(cre)Wwis^/Mmucd, MMRRC_015968-UCD) mice from MMRRC at UC Davis. *NeuroD1-GFP* (STOCK Tg (Neurod1-EGFP) CR99Gsat/Mmucd, MMRRC_000329-UCD) mouse strain was also obtained from MMRRC. These lines were maintained on a predominantly C57BL/6 J mixed background, with mice of both sexes used for experiments. Animals were housed under a 12 h light/dark cycle at 23 ± 3°C/50% humidity. Genotyping was performed at P4 for primary cultures and some immunostaining, and at around P30 for behavioral tests, RNA sequencing, and histological analyses.

### Genotyping

Tail or nail clippings were digested in 50 mM NaOH at 95°C for 10 min. After neutralizing by adding 1 M Tris-HCl (pH 8.0), samples were centrifuged at 4°C for 15 min. The supernatant was subjected for PCR using the following primers: Lig4^flox^ allele, 5’ GAGCTGCAACAGTTTGTGAAGTTTGTGAGGA 3’ and 5’ GTGTTGGTCAGGACCAGAAGGAAAGCA 3’; NeuroD1-Cre^cre^ allele, 5’ TAGGATTAGGGAGAGGGAGCTGAA 3’and 5’ CGGCAAACGGACAGAAGC 3’; Neurod1-GFP allele, 5’ TAGGATTAGGGAGAGGGAGCTGAA 3’ and 5’; Gabra6-Cre^cre^ allele, 5’ GATCTCCGGTATTGAAACTCCAGC 3’and 5’ GCTAAACAT GCTTCATCGTCGG 3’.

### In vivo and in utero electroporation and brain slice imaging

Plasmid electroporation into P5 cerebella was conducted using an electroporator (CUY21, Nepagene). A total of six square-wave pulses (70 mV, 50 ms duration, 150 ms intervals) were applied. Cerebella were harvested at P7 and embedded in 3.5% low-melting-point agarose (Nacalai, 01651-76), and coronally sliced into 300 μm-thick sections using a vibratome (NLS-AT, Dosaka EM). Slices were placed on Millicell-CM membrane inserts (Millipore, PICM0RG50) and embedded in a collagen gel matrix (FUJIFILM Wako, 631-00651). Slices were maintained in the medium composed of 60% basal medium Eagle (BME, Sigma-Aldrich, B9638), 25% Earle’s balanced salt solution (Sigma-Aldrich, E7510), 15% heat-inactivated horse serum (HS, Invitrogen, 26050070), 3 mM L-glutamine (Gibco, 35050-061), 1 mM sodium pyruvate (Sigma-Aldrich, S8636), 5.6 g/L glucose (Sigma-Aldrich, G7021), 1.8 g/L sodium bicarbonate (Sigma-Aldrich, S5761), and N-2 supplement (Invitrogen, 17502001). In utero electroporation into cerebral cortex was performed at E12. Brains were dissected at E14 and coronally sliced using a surgical knife^64^. Slices were mounted in a collagen gel matrix (FUJIFILM Wako, 631-00651) in DMEM/F12 (Sigma-Aldrich, D2906) supplemented with 5% HS, 5% fetal bovine serum (FBS, BioWest, S1400-500), penicillin/streptomycin (P/S, Invitrogen, 15140122), and N-2 supplement. Slices were placed in a stage-top incubator at 37°C with humidified 85% O₂/5% CO₂ gas flow, mounted on an upright microscope (BX61WI, Olympus)^25^. Time-lapse imaging was performed using a laser-scanning confocal microscope (FV1000; Olympus) equipped with a GaAsP detector and a 60× water-immersion objective lens (NA 1.1). Images were acquired every 10 min over 6 h.

### Live cell imaging on micropatterned substrates

The micropatterned PDMS substrate were made as previously described^27^. CGNs from P6 mouse cerebella were dissociated using the Neuron Dissociation Kit (Wako Pure Chemical Industries, 291-78001). For transfection, Nepa21-S (Nepagene) was used according to the manufacture’s instruction with a total of 6 μg plasmid DNAs. Following electroporation, cells were recovered in non-coated dishes for 2 h in 10% HS/BME, and then seeded on micropatterned PDMS substrates coated with laminin (Sigma-Aldrich, L2020) and incubated in culture medium BME with 26.4 mM glucose, 25 mM sodium bicarbonate, 1% bovine serum albumin (Sigma-Aldrich, A3156), 1x N-2 supplement}.

HeLa cells were maintained at ∼90% confluence in Dulbecco’s modified Eagle’s medium supplemented (DMEM, Invitrogen, 11965092) with 10% FBS and P/S on surface-treated culture dishes (Corning, 300-035). HeLa cells were transfected using Lipofectamine 2000 (Thermo Fisher Scientific, 11668019) one day after passaging and transferred to micropatterned substrates. Time-lapse imaging was performed using FV1000 with a 100× oil-immersion objective (NA 1.3) at 37°C in a humidified chamber with 5% CO₂. Images were captured at 5-min intervals.

### Transwell assay

Dissociated CGNs were seeded at 1.7×10^5^ cell/cm^2^ onto polycarbonate membrane with pore sizes of 0.4 μm, 3 μm or 8 μm (Corning, 3401, 3402, 3414, 3412, 3422) coated with poly-D-lysine (PDL, Sigma-Aldrich, P6407) and laminin. Drugs or DMSO (Nacalai, 13445-74) were added to both sides of the membrane 2 h after seeding. Cells were allowed to transmigrate for indicated durations and then either fixed or collected for further analyses. For transwell assays with transfected CGNs, cells were first plated on PDL- and laminin-coated 2.5 cm culture dishes (Corning) in culture media. At 24 h after plating, cells were washed with pre-warmed PBS and dissociated using Accutase (Nacalai, 12679-54) at 37°C for 5–10 min. Cells were collected by centrifugation, resuspended in a fresh culture medium, and then reseeded onto transwell inserts coated with PDL and laminin and incubated for indicated durations. For HeLa cells, transfected cells were suspended in DMEM and transferred to 1.12 cm^2^ transwell inserts that were preincubated with the medium at 1 × 10⁵ cells/well. The bottom chamber was placed with DMEM supplemented with 10% FBS to promote migration.

### Immunofluorescence of transwell cultures

Cells seeded on transwell inserts were fixed with 4% paraformaldehyde (PFA, Nacalai, 02890-45) and permeabilized with 0.1% Triton X-100 (Nacalai, 35501-02) in phosphate buffer saline (PBS) at RT. After blocking in 5% skim milk (Becton, Dickinson and Company, 232100), cells were incubated with primary antibodies diluted in blocking buffer overnight at 4°C. After thorough washes, cells were incubated with Alexa Fluor-conjugated secondary antibodies (Thermo Fisher Scientific) for 8–10 h at 4°C, followed by nuclear counterstaining with 4’-6-diamidino-2-phenylindole (DAPI; FUJIFILM Wako, 340-07971) (5μg/ml). The TUNEL assay was performed according to the manufacturer’s instructions for the Click-iT™ TUNEL Alexa Fluor™ Imaging Assay kit (Thermo Fisher Scientific, C10245). Membranes were then mounted using ProLong™ Gold Antifade Mountant (Invitrogen, P36961). Imaging was performed using a laser-scanning confocal microscope (FV1000-BX61, Olympus) through 20× (N.A. 0.5), 40× (N.A. 0.95) objectives or a 100× (N.A. 1.4) oil immersion objective, or a lattice structured illumination (SIM) microscope (Elyra 7, Zeiss) through a 63× (N.A. 1.40) oil-immersion objective.

### Immunofluorescence of brain tissues

Mice were anesthetized with isoflurane (FUJIFILM Wako, 099-06571) and perfused with 4% PFA in phosphate buffer (PB). Brains were dissected immediately and post-fixed in 4% PFA/PB for 4–6 h (for P15 or younger mice) or overnight (for P30 or older mice) at 4°C. Brains were washed in PBS then dehydrated in 30% sucrose (Nacalai, 30404-45) in PBS overnight at 4°C and were embedded in O.C.T. compound (Sakura Finetek,4583). Sagittal sections at 15 μm thickness were prepared using a cryostat (Leica, CM1950). For immunohistochemistry, sections were incubated with HistoVT One (Nacalai, 06380-05) at 70°C for 20 min for antigen retrieval. Following permeabilization with 0.5% Triton X-100/PBS and blocking, immunofluorescence was done as described above. For γH2AX immunostaining, all solution, including blocking buffer and antibody diluents, were prepared using Tris-buffered saline (TBS) instead of PBS. After staining with DAPI (10 μg/ml), sections were mounted using Fluoromount-G (Cosmo Bio, SBA-0100-01-25). Imaging was performed using the confocal and SIM microscope systems described above. For hematoxylin/eosin (HE) staining, anesthetized mice were perfused with PBS followed by 10% Formaldehyde Neutral Buffer Solution (Nacalai, 37152-51). Brains were post-fixed in the same fixative for 1 week at 4°C. Fixed tissues were processed for paraffin embedding using standard dehydration and clearing protocols, then sectioned at a thickness of 5 μm and subjected to HE staining. Images were acquired using an upright optical microscope (DM5000B; Leica) equipped with 2.5× (NA 0.07) and 10× (NA 0.3) objective lenses, an AdvanCam-U3X camera (Advan Vision), and AdvanView imaging software.

### TOP2**β**-DNA cleavage complex assay

Dissociated CGNs were seeded on transwell inserts of indicated pore sizes. After incubation for 8 or 12 h, cells either on the bottom of 3 µm or top of 0.4 µm membranes were washed once with pre-warmed (37°C) PBS and detached with Accutase at 37°C for 15 min. Cells were then rinsed with 10% FBS in culture medium, centrifuged at 4°C, and washed three times with cold PBS. TOP2 binding assay was performed as previously described with a few modifications^34^. Briefly, cell pellets were resuspended in 1% N-Lauroylsarcosine Sodium Salt in TE buffer (diluted from 30% Sarcosyl NL-30, Nacalai, 20135-14). A total of 2 ml of lysate was loaded onto a density-gradient cushion of cesium chloride (Nacalai, 07807-11), consisting of layers with densities of 1.86, 1.7, 1.5, and 1.45 g/ml, from bottom to top. Sample were subjected to ultracentrifugation at 100,000 × g for 16 h at RT. One ml aliquots were collected from the top to the bottom of the gradient. For slot blot analysis, 100 µl of each aliquot was loaded to a PVDF membrane pre-treated with methanol using a Bio-Dot apparatus (Bio-Rad, 1706545). After washing with 0.2 M PB, the membrane was incubated overnight with an anti-TOP2β antibody in 5% skim milk. The membrane was then incubated with a horseradish peroxidase-conjugated anti-rabbit secondary antibody. Image Gauge software (Fujifilm) was used to detect TOP2β signals.

### AAV Injection

AAVs (10⁹–10¹⁰ plaque-forming units) were produced using AVB Sepharose High Performance (GE Healthcare, 28-4112-01) following the manufacturer’s instructions. Briefly, transfected 293T cells were harvested, followed by freeze-thaw cycles (three times using liquid nitrogen and a 37 °C water bath). Cell lysates were treated with 5-10 µl Benzonase, incubated at RT for 5 min, and centrifuged. The supernatant was filtered before bead binding. 400 µl AVB beads were washed three times with washing buffer (20 mM Tris-Cl pH 8.0, 250 mM NaCl, 10 mM MgCl₂) and incubated with the viral lysate at room temperature for 15 min. Beads were then washed three times and transferred to a column Mobicol “Classic” (MoBiTec, M1002). Bound AAVs were eluted acidic elution buffer (250 mM NaCl, 10 mM MgCl₂, pH 3.0 adjusted with HCl) and immediately neutralized with 1 M Tris-HCl (pH 8.0). Eluted virus was concentrated to 100–200 μl using Amicon Ultra-4 Ultracel 50K centrifugal filters (Millipore, UFC805024).

Mice at the age of 12 mo were anesthetized with an intraperitoneal injection (0.1 ml/10 g body weight) of a mixed anesthetic solution containing 0.03 mg/ml medetomidine (Kyoritsu Seiyaku), 0.4 mg/ml midazolam (Astellas Pharma Inc), and 0.5 mg/ml butorphanol (Meiji Seika Pharma). A total of 1 µl of virus was injected over 1 min at a stereotaxic location 6.84 mm posterior to bregma, along the midline, and 1.5 mm below the dural surface. The needle was left in place for an additional 5 min post -injection before being slowly withdrawn. Following injection, the incision was sutured, and mice received an intraperitoneal injection of 0.3 mg/ml atipamezole (Kyoritsu Seiyaku) for anesthesia reversal. Animals were kept in a warm (37°C) recovery cage until fully awake. One week after injection, mice were perfusion-fixed. Brains were post-fixed, washed in PBS, and embedded in 4% low-melting-point agarose. Coronal sections of 100 µm thickness were prepared using a vibratome Immunostaining for GFP and vglut1/2 was performed as described. Images were acquired using a laser-scanning microscope (Andor Dragonfly 500) equipped with a 100× oil immersion objective (NA 1.49).

### Electron microscopy

Brains fixed in 4% PFA/PB were washed in PB followed by distilled water. First post - fixation was carried out on ice using 2% osmium tetroxide (OsO₄) with 1.5% potassium ferrocyanide for 2 h. After rinsing, second post-fixation was performed at room temperature using 1% OsO₄ in PB for 2 h. Tissues were then dehydrated through graded ethanol (50–100%), cleared in propylene oxide, and infiltrated with EPON resin through graded propylene oxide: EPON mixtures, followed by overnight incubation in 100% EPON. Polymerization was conducted at 45°C for one night and 60°C for two nights.

Semi-thin sections (700 nm–1 μm) were prepared for light microscopic screening, and ultrathin sections (60–80 nm) were used for electron microscopy. Sections were stained with uranyl acetate and lead citrate, and images were captured using a transmission electron microscope (Hitachi, H-7650 or JEOL, JEM-1400Flash).

### RNA-seq

Total RNA was extracted from: dissociated CGNs from wildtype P6 ICR mice cultured for 8 h on normal dishes and transwell inserts; control and *Lig4 ^ND^*^1^*^-Cre^* cerebella at 2 mo using the miRNeasy Mini Kit (QIAGEN, 217004) and assessed for quality using a BioAnalyzer (Agilent Technologies). Libraries were prepared using the TruSeq Stranded mRNA Library Prep Kit (Illumina) and sequenced on the NovaSeq 6000 system (Illumina) with 100 bp single-end reads. Raw Fastq files were trimmed and aligned to the mouse reference genome (mm10) using STAR (v2.7.11a)^65^. Mapped reads were assembled with FeatureCounts (v2.0.8)^66^, and differential gene expression was analyzed using DESeq2(v2.11.40.8)^67^ based on read counts. GO analysis was performed using upregulated and downregulated DEGs(p < 0.05, determined by DESeq2) via the DAVID functional annotation tool^68,69^.

For public data analysis, raw sequence files were obtained from the BioProject database (PRJNA281127)^42^ and from the GEO database (GSE212336^43^, GSE221124^44^, and GSE174265^45^). The reads were mapped to mouse genome (mm10) using hisat2 (v2.1.0)^70^, and mapped reads were assembled with FeatureCounts (v2.0.0).

### ChIP-seq

Dissociated CGNs cultured on PDL and laminin-coated dishes (Corning) were collected with Accutase as described above. For tissue preparation, cerebella were dissected from P15 mice. Nuclei were isolated according to a previously reported protocol^71^. The nuclei pellets were resuspended and filtered through a 30 μm cell strainer (Miltenyi Biotec, 130-098-458) into pre-chilled tubes. ChIP for H3K4me1, H3K27ac, H3K4me3, H3K9me3, and LaminB1 was performed as previously published methods^40^. Briefly, isolated nuclei or cell pellets were resuspended in pre-warmed medium and fixed with 1% formaldehyde (Sigma-Aldrich, F1635) at 37°C for 10 min. Fixation was quenched by adding 1 M glycine/PBS to a final concentration of 125 mM. Cells were washed twice with cold PBS. For lysis, fixed cell pellets were resuspended in cold RIPA buffer (10 mM Tris-HCl pH 7.5, 1 mM EDTA, 0.1% SDS, 0.1% sodium deoxycholate, 1% Triton X-100) supplemented with protease inhibitors (ThermoFisher, 78442). Chromatin was sheared using a Covaris S220 sonicator (20% duty cycle, 175 peak powers, 200 cycles/burst) for 30 min at 4°C. Lysates were cleared by centrifugation and precleared with Dynabeads Protein A (Invitrogen, DB10002) for 30 min at 4°C. Antibodies (10 μg) were conjugated to pre-washed Dynabeads in PBS for 10 min at room temperature, washed, and incubated with chromatin overnight at 4°C with rotation. Beads were then sequentially washed: twice with RIPA buffer, twice with RIPA + 0.3 M NaCl, twice with LiCl buffer (0.25 M LiCl, 0.5% Igepal-630, 0.5% sodium deoxycholate), once with TE + 0.2% Triton X-100, and once with TE. Crosslinks were reversed by incubation at 65°C for >4 h in 0.3% SDS and 1 mg/ml Proteinase K (Nacalai, 15679-06). DNA was purified and eluted in 10 mM Tris-HCl. DNA obtained from ChIP was used to construct sequencing libraries using the KAPA HyperPrep Kit (KAPA Biosystems). Libraries were sequenced on the NextSeq 500 platform (Illumina) using 75 bp single-end reads.

### ATAC-seq

Dissociated CGNs cultured on PDL and laminin-coated dishes (Corning) were collected with Accutase as described above. Cells were resuspended in ice-cold lysis buffer (10 mM Tris-HCl pH 7.4, 10 mM NaCl, 3 mM MgCl₂, 0.1% IGEPAL CA-630) and centrifuged again under the same conditions. The supernatant was removed. For transposition, 2× Tagmentation Buffer (Diagenode, C01019043) and Tagmetases (Diagenode, C01070012-10) were diluted in water to make reaction mixture. The reaction mixture was then added to cell pellet and incubated at 37 °C for 30 minutes. DNA was purified using the QIAGEN PCR Purification Kit (QIAGEN, 28104) and eluted in elution buffer. Purified ATAC-DNA was used to library amplification with Q5 Hot Start High-Fidelity 2× Master Mix (NEB, M0494S). Libraries were sequenced on the NextSeq 500 platform (Illumina) using 75 bp single-end reads.

### END-seq

Dissociated CGNs were seeded either on 3-μm transwell inserts or on culture dishes and incubated for 6 h in the presence of Palbociclib (5 µM). To stabilize DNA breaks, both control and migration conditions were treated with etoposide (50 μM) at 3 hours after seeding to ensure sufficient cell attachment. Cells were collected from the bottom side of the transwell inserts or directly from culture dishes with Accutase. For tissue samples, nuclei were isolated from P15 control and *Lig4 ^ND^*^1^*^-Cre^* cerebella as described above.

Isolated nuclei or cells were embedded in agarose plugs and END-seq was performed as previously described^39^. Agarose plugs were treated with proteinase K for 1 hour at 50°C and then for 7 hours at 37°C. Etoposide was maintained at 50μM in all the steps until digestion with Proteinase K. Plugs were washed in wash buffer 10 mM Tris-HCl pH 8.0, 50 mM EDTA) and in TE (10 mM Tris-HCl pH 8.0, 1 mM EDTA) followed by RNase A treatment for 1 hour at 37°C. Inside of the plugs, DNA ends were blunted for 1 hour at 37°C with Exonuclease VII followed by Exonuclease T (NEB) for 1 hour at 25°C to detect TOP2 breaks. After blunting, A-tailing was performed, followed by biotinylated END-seq hairpin adaptor 1 ligation using NEB Quick Ligase. Agarose plugs were melted and dialyzed and DNA was sonicated using Covaris S220 sonicator for 4 min at 10% duty cycle, peak incident power 175, 200 cycles per burst, at 4°C. DNA was ethanol-precipitated and dissolved in 80 μl TE buffer. Biotinylated DNA was isolated using MyOne Streptavidin C1 Beads (ThermoFisher, 650–01), followed by end repair (dNTPs, T4 polymerase (NEB), Klenow (NEB), T4 PNK) and dA-tailing (Klenow exo- (NEB), dATP). The second end was ligated to “END-seq hairpin adaptor 2” using NEB Quick Ligase. Hairpins were digested using USER (NEB), and the resulting DNA fragments were PCR amplified using Illumina double barcoded primers. PCR fragments were isolated by size selection from agarose gel and purification using NEB Monarch Gel Extraction Kit. Libraries were quantified using KAPA Library Quantification Kit and sequenced using Illumina NextSeq 550 and NovaSeq X.

### Genomic analysis

END-seq, ATAC-seq and ChIP-seq reads were mapped to the mouse (GRCm38p2/mm10) genomes using Bowtie2 (v2.5.1-1)^72^ with the default parameters. For END-seq, peaks were called using MACS(v1.4.3)^73^ with the parameters:-nolambda,-nomodel and-keep-dup = all (keep all redundant reads) and subsequent analysis were done using bedtools (v2.31.1)^74^ and R (v4.3.2). For ETO treated samples we used the corresponding non-treated samples as control, keeping > 5 fold-enriched peaks. Peaks were later filtered by >1.5 RPKM and >3 fold RPKM to the control. Condition or genotype -specific peaks were selected as not called in the other condition/genotype and more than 2 fold RPKM. Common peaks were defined as filtered peaks that overlapped between conditions/genotypes.

The UCSC database^75^ was used to obtain the RepeatMasker annotations and Transcription start sites (TSS), transcription end sites (TES), exons, and introns positions (RefSeq annotation table). H3K9me3, H3K4me3, H3K4me1, H3K27ac ChIP-seq peaks were called using epic2^76^(with -bin 10000 for H3K9me3 and 500 for the rest). Lamin B1 domains (regarded as LADs) were called using Enriched domain detector (Edd)^77^ with --bin-size 37 and -g 37. ATAC-seq peaks were called using Genrich default parameters, available at https://github.com/jsh58/Genrich. RNA-seq, ATAC-seq and histone marks ChIP-seq peaks in control samples were used to define chromatin categories (active promoters, active enhancers, active exons and introns, transcriptionally inactive and unlabeled euchromatin, heterochromatin with LADs and heterochromatin without LADs) using bedtools as follows, “active promoters”: H3K4me3 peaks that overlap with ATAC-seq peaks; “active enhancers”: ATAC-seq peaks that do not overlap with H3K4me3; “active exons” exons of genes in RNA-seq with DESeq2 Base Mean value >=5 after subtracting H3K4me3 and ATAC-seq peaks; “active introns”: introns of genes in RNA-seq with DESeq2 Base Mean value >=5 after subtracting H3K4me3 and ATAC-seq peaks; “heterochromatin LADs”: LaminB1 LADS after subtracting the union of active promoters, active enhancers, active exons, active introns; “heterochromatin noLADs”: H3K9me3 peaks after subtracting LaminB1 LADS, active promoters, active enhancers, active exons and active introns”; “Euchromatin”: genome mm10 after subtracting heterochromatin; “transcriptionally inactive and unlabeled euchromatin”: euchromatin after subtracting active promoters, active enhancers, active exons and active introns.

Chromatin categories in cerebellar cells from P15 control and *Lig4 ^ND^*^1^*^-Cre^* mice were defined using bedtools as follows, “active promoters”: H3K4me3 peaks that overlap with H3K27ac peaks; “active enhancers”: H3K4me1 peaks that overlap with H3K27ac peaks after subtracting active promoters; “poised enhancers”: H3K27ac peaks after subtracting active promoters and active enhancers; “heterochromatin”: H3K9me3 peaks; “euchromatin”: genome mm10 after subtracting heterochromatin; “transcriptionally inactive and unlabeled euchromatin”: euchromatin after subtracting active promoters, active enhancers and poised enhancers.

CTCF and RAD21 ChIP-seq peaks were called using MACS (v1.4.3) with default parameters. Position Weight Matrix (PWM) of CTCF was taken from JASPAR database^78^ and significant CTCF motifs (p < 1×10^−4^) in the mouse genome were selected using FIMO tool^79^. Significant CTCF motifs at double CTCF/RAD21-bound peaks were used for the analysis. For data visualization bedgraph generated with bedtools were converted to bigwig files using bedGraphToBigWig^80^. Visualization of genomic profiles was done by the UCSC browser^81^. Genome browser profiles were normalized to present RPM. For aggregate plots around CTCF sites, TSSs and gene bodies signal was smoothed using smooth.spline function in R.

### Behavioral Tests

All behavioral tests were conducted during the light phase in the room where the mice were kept. Age- and weight-matched male mice were used for all behavioral tests.

#### Footprint analysis

Each mouse was guided to walk along a straight runway lined with paper toward a dark box at the end. The forepaws and hindpaws were coated with non-toxic black and red paints, respectively. Each mouse completed 3 consecutive trials. A fresh sheet of white paper was placed on the floor of the runway for each run. Parameters measured included stride length, width based on forelimb or hindlimb placement, and the distance between the same-side forepaw and hindpaw prints (gait overlap)^82^.

#### Balance beam test

Mice were encouraged to cross a narrow cylindrical plastic beam (12 mm diameter, 1 m length), placed horizontally 1 m above the bench surface, toward a dark goal box. Each mouse was trained for the task for 2 days before trial experiments. Mice performed 3 trials per day with a 5-min interval between trials. Mice were allowed to cross the beam voluntarily without external stimuli during trials. Hindlimb slips were recorded only while passing through the middle 80 cm of the beam. The average number of slips from 3 trails was calculated for each mouse.

#### Rotarod test

Mice were trained to remain on a rod rotating (Rota-rod treadmill, Muromachi Kikai, MK-600) at a constant speed of 2 rpm for 180 sec^83^. After a 1 h interval, a 180-sec test session was conducted on a rotating rod with acceleration from 2 rpm to 20 rpm. 1 trial were run for each mouse per day for 3 consecutive days, with the training session performed only on Day 1. The latency to fall (time spent on the rod) was recorded for each mouse in each session.

#### Grip strength measurement

Grip strength was assessed using a grip strength meter (Force gauge. A and D, AD-4932A-50N) with a rectangular grid handle. Mice were allowed to grasp the mesh of the grid handle with all four limbs. Once they had a firm then they were slowly pulled horizontally by the tail until all limbs released the grid. The maximum force exerted during the trial was recorded. 5 trials were performed for each mouse, with 5 min rest period between trials. The maximum force recorded from the 5 trials was used for analysis. Grip strength was normalized by calculating the ratio of grip strength to body weight, expressed a raw grip strength (newton)/weight (g) × 100.

### Image analysis and data representation

All image processing and analysis were performed using FIJI (ImageJ), except for the images presented in supplementary Fig. 3b, which were processed with Imaris (Oxford Instruments). For fluorescence images, maximum projections of multiple z stacks were used in all figures except the magnified viewed enlarged in the supplementary fig. 2f, a single Z stack was used to show the mono-structure of γH2AX. The γH2AX-positive ratio in Fig. 1c was quantified based on P27kip1-positive cells, excluding those in the outer EGL, which was quantified based on the number of DAPI-stained cells. StarDist-0.3.0, a tool plug-in on FIJI, was used to count the number of nuclei. In transwell assays, cells with more than five nuclear γH2AX foci, and cells with one or more nuclear 53BP1 foci were counted as positive.

### Statistical analysis

Statistical analysis performed by GraphPad Prism 7 or R.4.3.3. Details regarding the number of biological replicates are provided in the respective figure legends. Error bars in the graphs indicate standard deviation. The statistical methods applied are specified within each figure legend. Statistical significance is represented as follows: *p < 0.05; **p < 0.01; ***p < 0.001.

## Acknowledgement

We thank F. Alt for kindly providing the *Lig4 ^flox/flox^* mice; S. Kawaguchi and T. Inoshita for advice and assistance with rotarod test; T. Ogi and S. Saitoh and their group members, M. Piel, A. Baffet, S. Takeda, M. Foiani, and L. Nguyen for advice and discussion; Division of Electron Microscopic Study, Center for Anatomical Studies, Medical School of Kyoto Univ. for technical assistance with electron microscopy and histological analyses; S. Yawata and L. Peng for advice and assistance with AAV injection. This work was supported by AMED 23gm1910002, grants from Japan Society for the Promotion of Science (JSPS) Kakenhi (22H05169, 21H00250 and 21K19312), Uehara Memorial Foundation, Takeda Science Foundation to MK.

## Author Contributions

Z.Z. and M.K. conceived and designed the study and wrote the manuscript. Z.Z., P.Z., N.T., T.K., N.N., M.U. and J.K. performed migration assays and histological analyses of wildtype and mutant mice. Z.Z., P.Z. and N.T. performed behavioral analyses. Z.Z., Y.K., M.S. and A.C. performed RNA-seq, ATAC-seq, ChIP-seq and END-seq analyses. T.F. assisted AAV-injection and performed electron microscopy experiments. F.I. assisted SIM imaging. H.S. and Z.Z. performed TOP2cc measurement.

The authors declare that they have no competing interests.

## Data availability

All of the data and reagents in this study are available from the corresponding author upon reasonable request.

