## Supplementary Figures for "Neuronal migration induces DNA damage in developing brain"

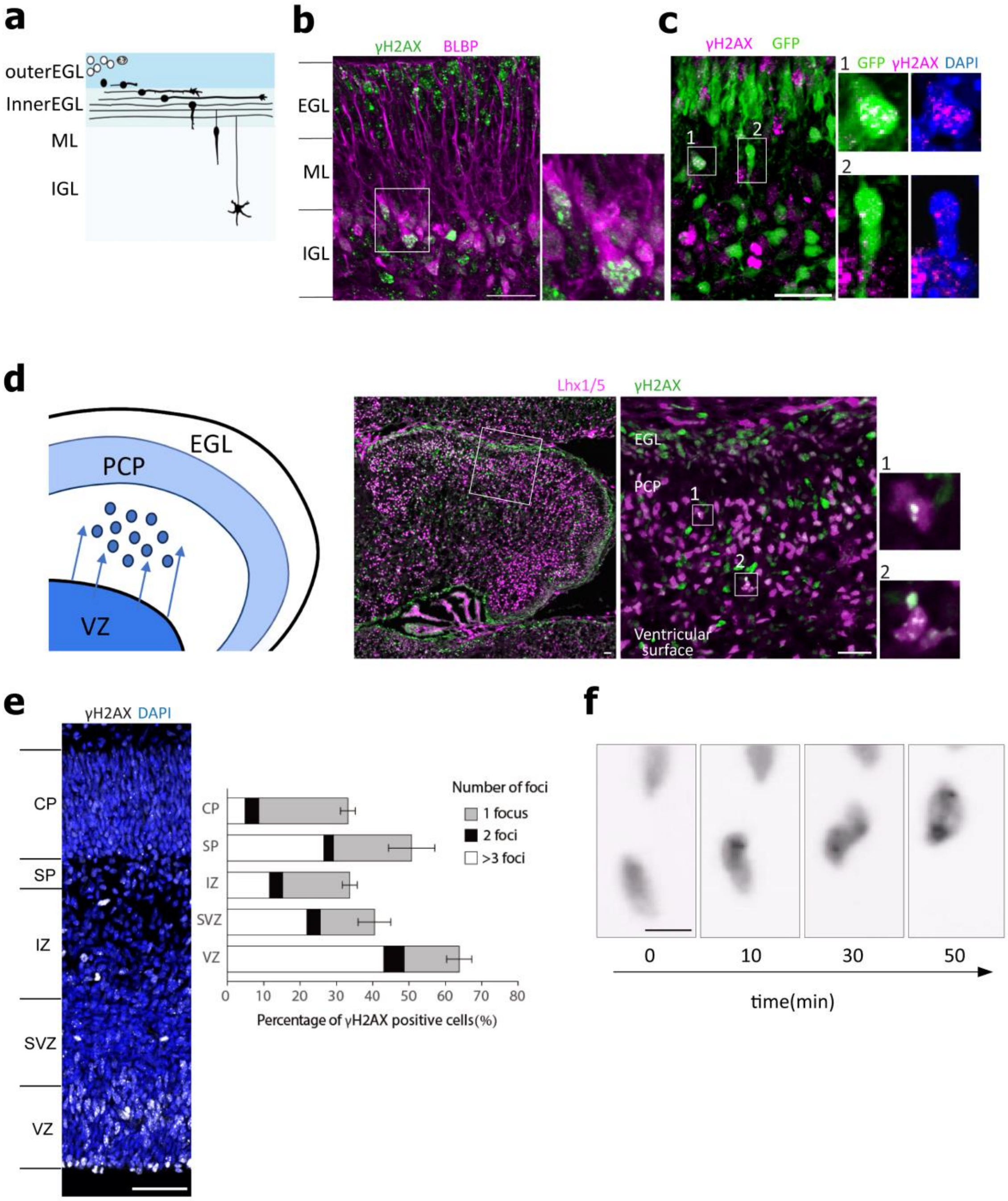


**Supplementary Fig. 1. DSB formation during migration is common across multiple types of postmitotic neurons.**

**a,** Schematic illustration of CGN differentiation during postnatal cerebellar development. CGPs (white) and CGNs (black) populate the outer and inner EGL, respectively. CGNs extend axons and migrate along the EGL surface, and then radially through the ML to reach the IGL. **b,** Sagittal section of P6 cerebellar cortex stained with γH2AX (green) and BLBP (magenta). Some γH2AXfoci in the IGL are seen in BLBP-positive glial cells. Right panel showed a magnified view of the boxed region. **c**, Sagittal section of the cerebellar cortex from a Neurod1-GFP mouse at P6 stained with γH2AX (magenta), GFP (green) and DAPI (blue). GFP-positive CGNs migrating in the ML are boxed and magnified on the right. γH2AX foci are seen in the nucleus. **d,** *Left,* Schematic of Purkinje cell migration in the cerebellar primordium during embryonic development. *Right 3 rows,* E17 cerebellum stained for γH2AX (green) and Lhx1/5 (magenta). The mid panel is an enlarged image of the boxed region of a low magnified view on the left. Some Lhx1/5-positive Purkinje cells migrating toward the Purkinje cell plate (PCP) bear γH2AX foci (boxes 1 and 2). **e,** *Left*, E15 cerebral cortex stained for γH2AX (white) and DAPI (blue). *Right*, The percentage of γH2AX-positive cells in each layer. Samples were collected from more than 3 independent experiments (n = 5 mice). **f,** Time-lapse images of a neocortical neuron migrating in the cortical plate in a slice culture prepared from E14 mouse brain. 53BP1-mNG (gray) was introduced by in utero electroporation at E14. CP, cortical plate; SP, subplate; IZ, intermediate zone; SVZ, subventricular zone; VZ, ventricular zone. Data represent mean ± s.d. Scale bars; **b-d**, 24 μm; **e**, 50 μm; **f,** 5 μm.

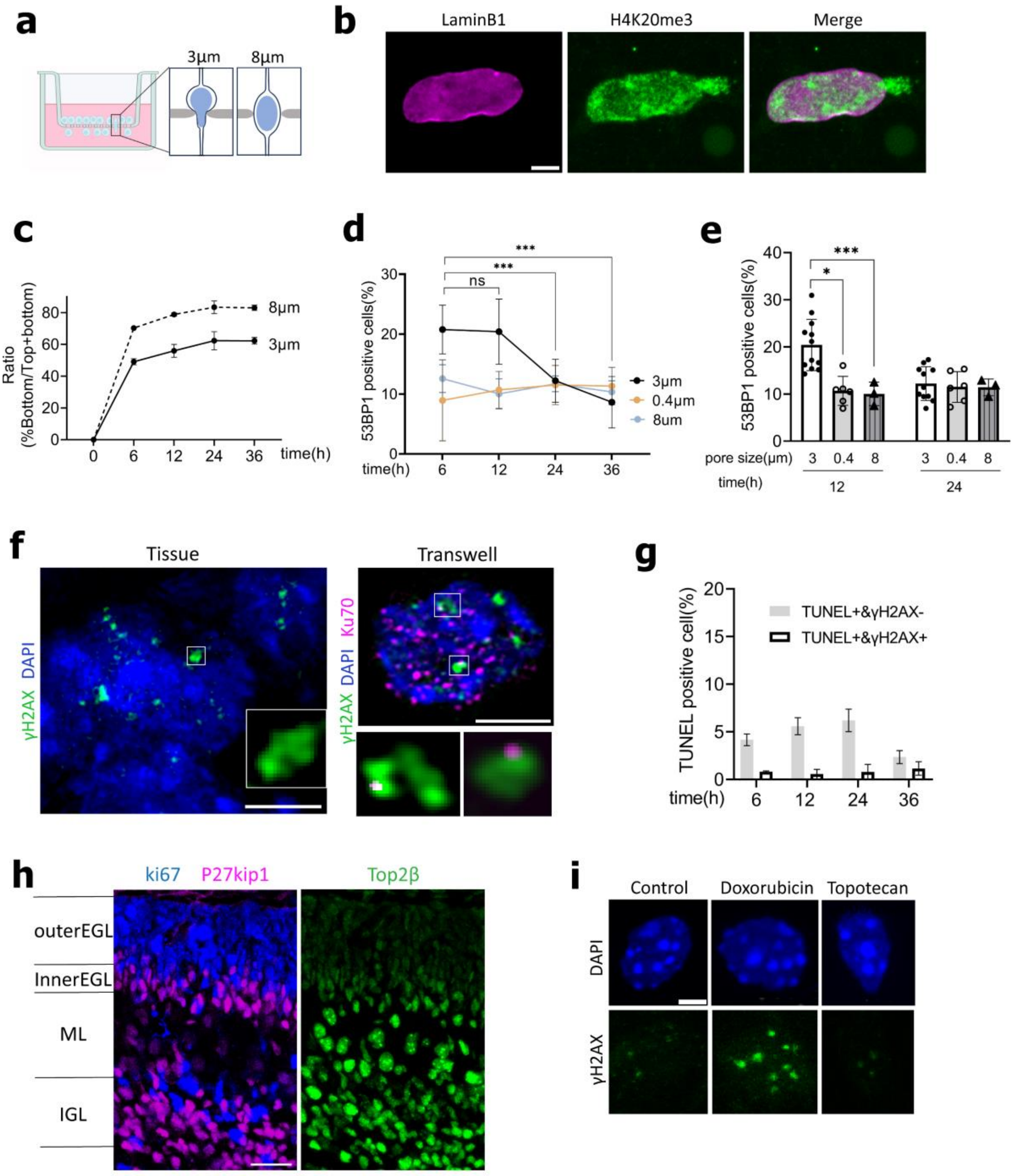


**Supplementary Fig. 2. CGNs generate DSBs during confined migration.**

**a,** Transwell assay. **b,** A CGN with nuclear blebbing after transwell migration through 3-μm pores. Chromatin (H4K20me3, green) is herniated from a Lamin B1-negative site (magenta). **c,** The percentage of CGNs that have completed transwell migration at indicated time after inoculation. Samples were collected from 4 independent experiments. **d, e**, The percentage of 53BP1-positive CGNs after transwell migration through confined (3-μm) and non-confined (8-μm) pores, as well as non-migrated cells on 0.4-μm pore filters, at different time points after seeding. Samples were collected from at least 3 independent experiments per group. **f,** Super-resolution images of CGNs in the IGL of P6 cerebellar cortex (left) and a cultured CGN migrated through 3-μm pores (right), immunostained for γH2AX (green) with or without Ku70 (magenta). γH2AX nanoclusters in the boxed regions are magnified. **g**, Quantification of TUNEL-positive cells after transwell migration through 3-μm pores at different time points after inoculation. Samples were collected from 3 independent experiments. **h,** Immunostaining of P6 cerebellar cortex showing TOP2β (green), Ki67 (blue), and P27kip1 (magenta) expression. **i,** CGNs migrated through 3-μm pores with or without Doxorubicin or Topotecan. Cells were fixed at 24 h after inoculation and stained for γH2AX. Scale bars; **b, f**, **i**, 3 μm; **h**, 24 μm. Data represent mean ± s.d. *p < 0.05, ***p < 0.001, two-way ANOVA with Tukey's multiple comparisons tests.

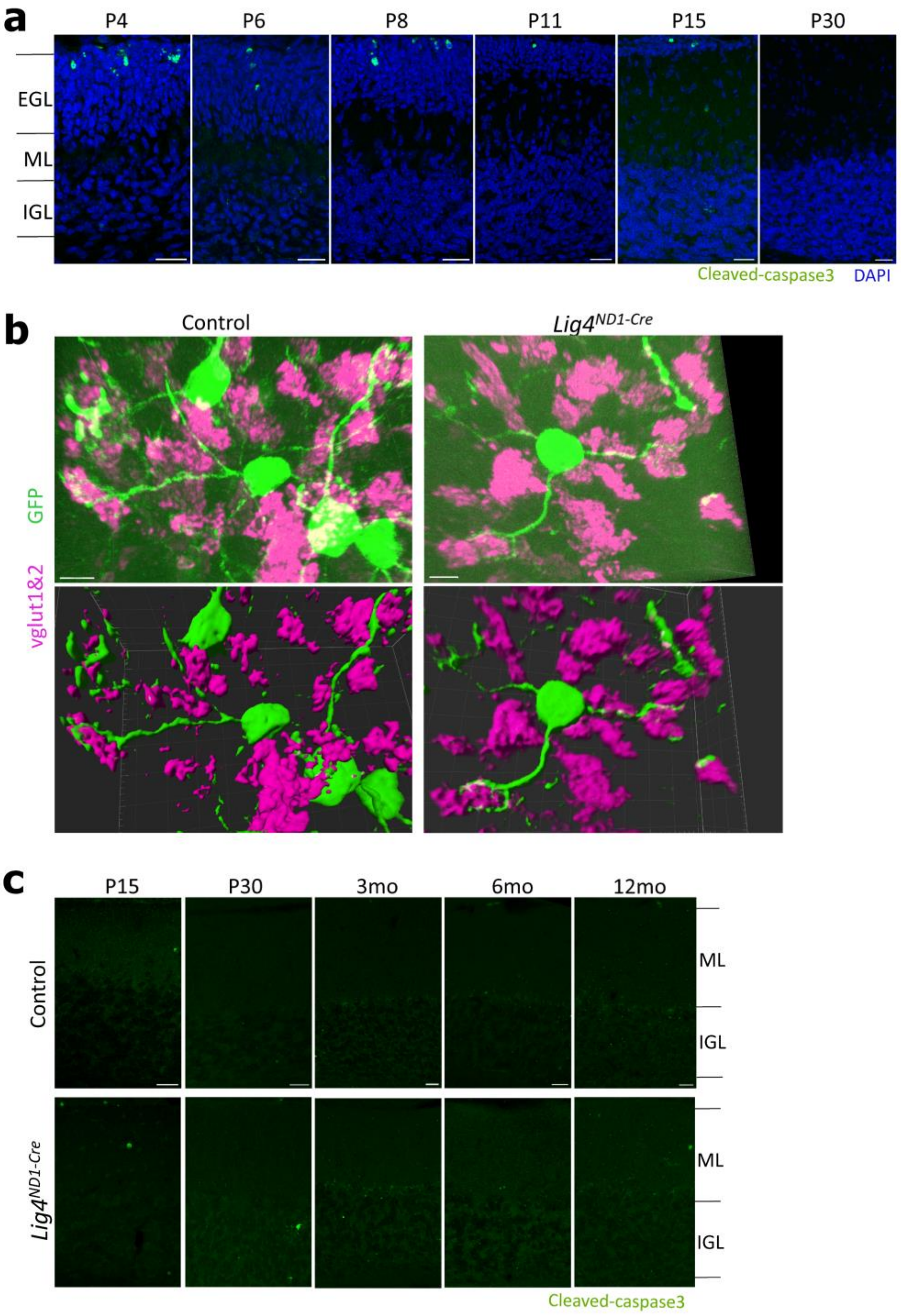


**Supplementary Fig. 3. Persistent DSBs in *Lig4*$^{ND1\text{-}Cre}$ do not induce cell death.**

**a,** Immunostaining of the developing cerebellar cortex with an apoptosis marker cleaved caspase-3. Signals are scattered in the outer EGL populated with mitotic CGPs, but are not seen in the inner EGL and IGL. **b,** 3D stack views (top) and surface rendering images (bottom) of CGNs (GFP, green) and mossy fiber terminals (vglut1/2, magenta) in control and *Lig4* $^{ND1\text{-}Cre}$ cerebellar cortices at 12 mo. CGNs were labeled by administrating AAV-GFP by injection at 12 mo and subjected for immunofluorescence. **c,** Immunostaining of the cerebellar cortices from control and *Lig4* $^{ND1\text{-}Cre}$ mice at different ages with cleaved caspase-3 (green). Scale bars; **a**, **c**, 24 µm; **b,** 5 µm.

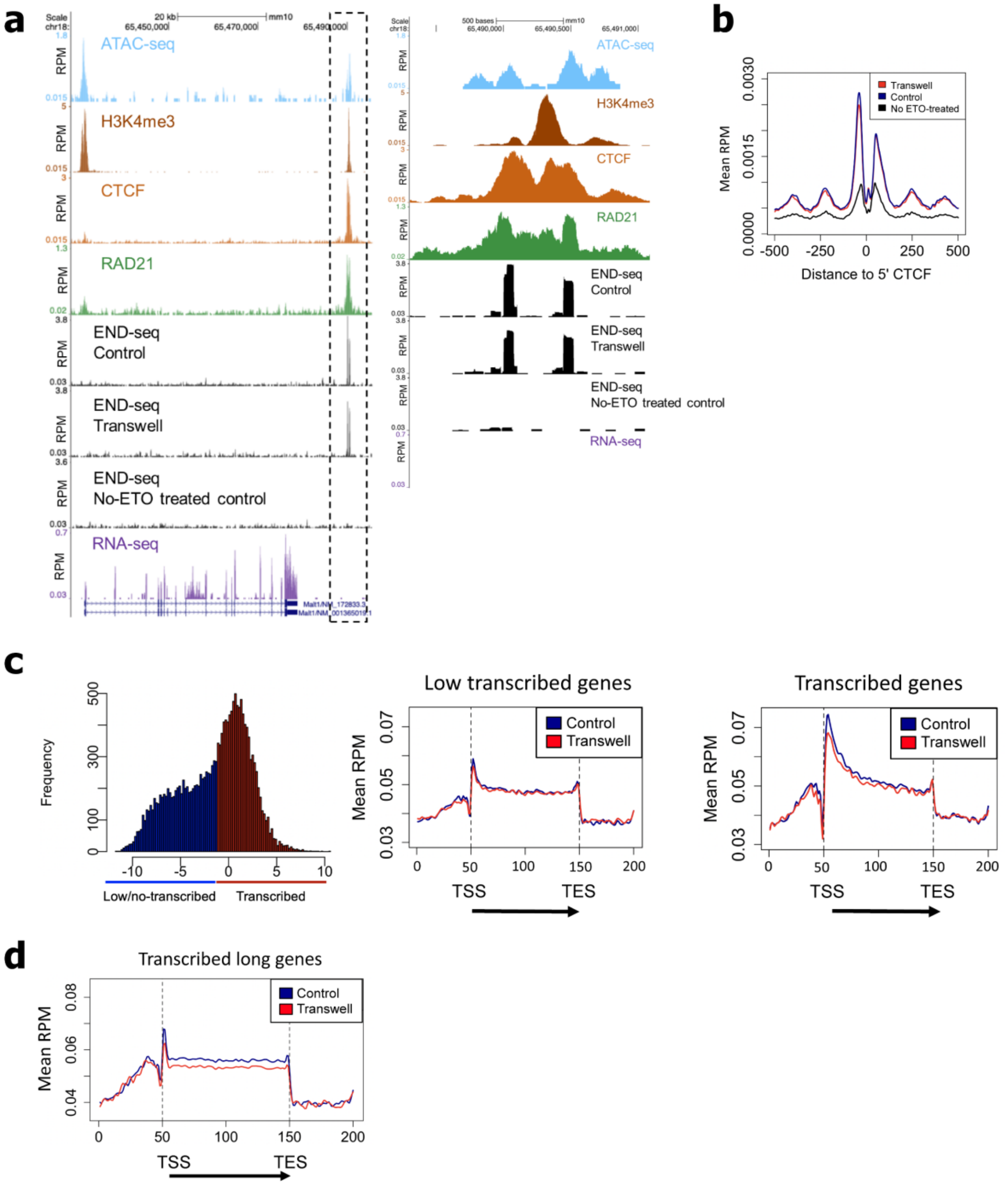


**Supplementary Fig. 4. Genome-wide mapping of mechanostress-induced DSBs in migrating CGNs.**

**a**, Genomic profile showing an example of etoposide-induced END-seq peaks in CGNs with or without confined migration in a site located at chr18:65428139-65495592. Boxed region in broken lines is enlarged on the right. From top to bottom: chromosome accessibility, measured by ATAC-seq; H3K4me3, CTCF and RAD21 occupancy measured by ChIP-seq, and transcription by RNA-seq, in unmigrated control CGNs; DSB profiles in etoposide-treated CGNs with or without transwell migration, and unmigrated CGNs without etoposide (NT). **b**, Aggregate plot of DSBs in migrated (red) and unmigrated (dark blue) CGNs ± 500 bp from the 5' end of the CTCF motif (position 0) at sites bound by both CTCF and RAD21. **c**, *left,* Histogram of transcription levels of genes assessed by RNA-seq in control CGNs. Colors indicate expression levels from low/non-transcribed (dark blue) to highly transcribed (red) based on the median of $\log_2$ (RPKM) value of all genes. Middle and right: Distribution of DSBs (reads per million mapped reads) in the gene bodies of low/non-transcribed genes (middle) and transcribed genes (right) in migrated and unmigrated CGNs. Gene bodies (TSS to TES) were divided in 100 equal-length bins, with 2,000 bp flanking regions upstream of the TSS and downstream TES binned in 50 intervals each; genes shorter than 2,000 bp were excluded. **d**, Distribution of DSBs in the gene bodies of neuronal long genes (the top 10% by gene lengths) that are transcribed in control CGNs, in migrated (red) and unmigrated (dark blue) CGNs.

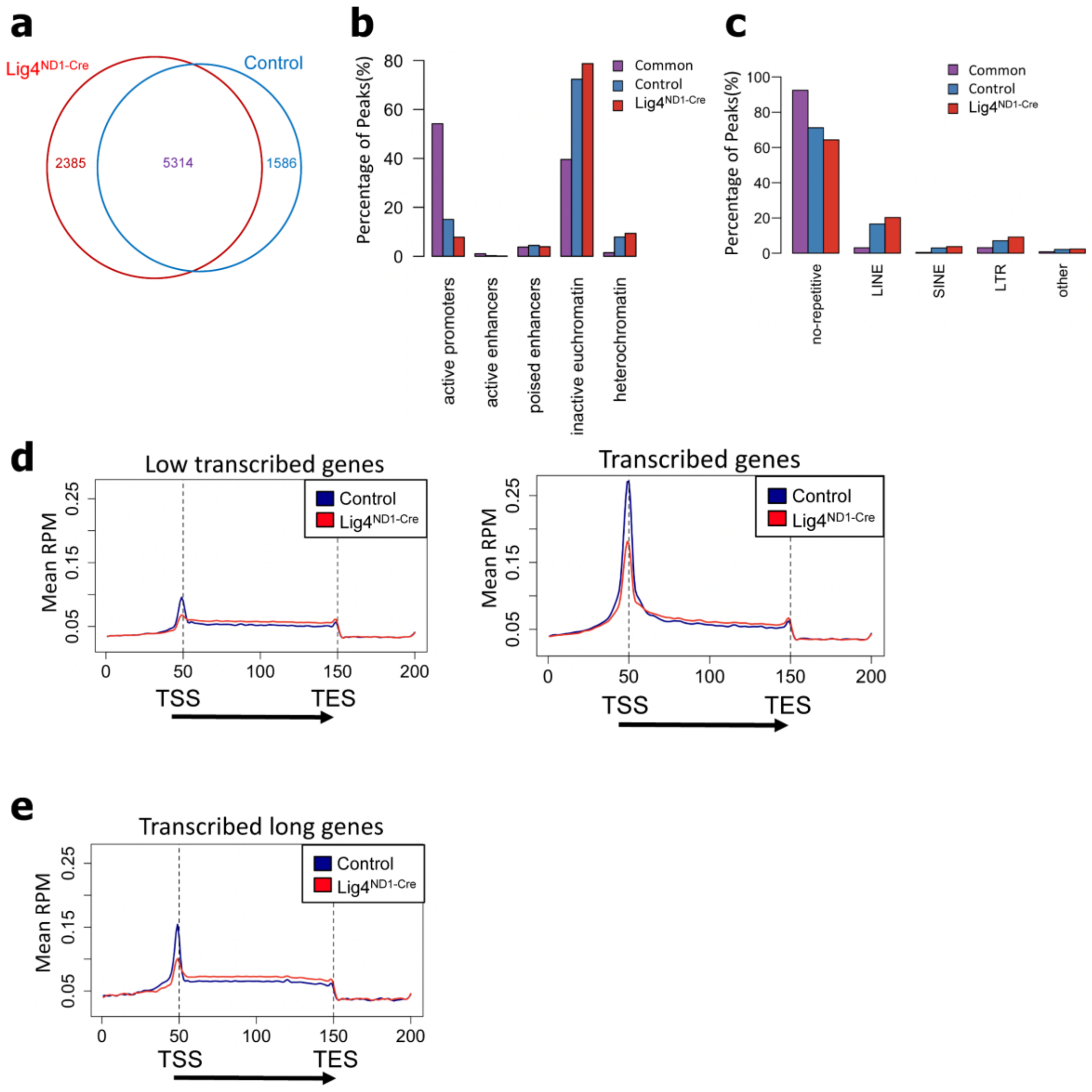


**Supplementary Fig. 5. Genome-wide mapping of persistent DSBs in *Lig4*$^{ND1-Cre}$ cerebellum.**

**a**, Venn diagram showing the overlap of END-seq peaks in the cerebellar cells from P15 control and *Lig4*$^{ND1-Cre}$ mice (without etoposide treatment). **b,** Differential END-seq peaks classified by chromatin categories (see Methods). **c**, Relative levels of END-seq peaks in repetitive elements. **d**, Distribution of DSBs in the gene bodies of low/non-transcribed (left) and transcribed genes (right) in cerebellar cells from control (dark blue) and *Lig4*$^{ND1-Cre}$ (red) mice. **e**, Distribution of DSBs in the gene bodies of long neuronal genes (the top 10% by gene lengths) in cerebellar cells from control (dark blue) and *Lig4* $^{ND1\text{-}Cre}$ (red) mice. Data showing one of two biologically independent replicates; similar results were obtained in the second.

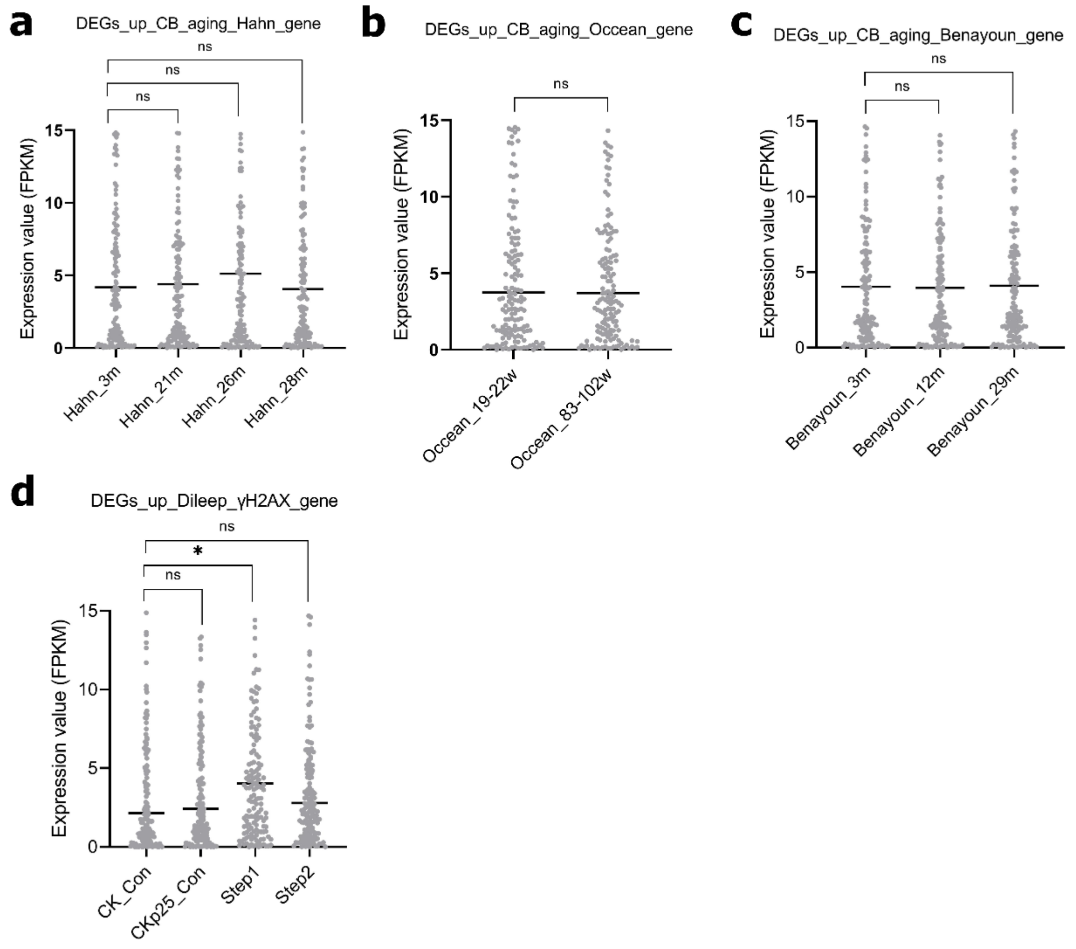


**Supplementary Fig. 6. Transcriptional changes in *Lig4* $^{ND1\text{-}Cre}$ compared to aged or DNA-damaged cerebra and cerebella.**

Quantification of expression values (FPKM) from published datasets for the list of upregulated DEGs (n = 170 genes in **a-c**; n=166 genes in **d**) generated by DEseq2 analysis from our RNA-seq datasets. **a-c,** Mean FPKM of individual genes in cerebellar samples from Hahn et al., 2023 (**a**)**,** Occean et al., 2024 (**b**) and Benayoun et al., 2019 (**c**). **d,** Mean FPKM of individual genes in cerebral samples from controls (CamKII promoter-tTA mouse and CK-p25 mice control) and neurodegeneration step 1 and 2 (Dileep et al., 2023). Horizontal lines represent median. *$p < 0.05$. **a, c, d,** Kruskal-Wallis test; **b,** Mann-Whitney test.

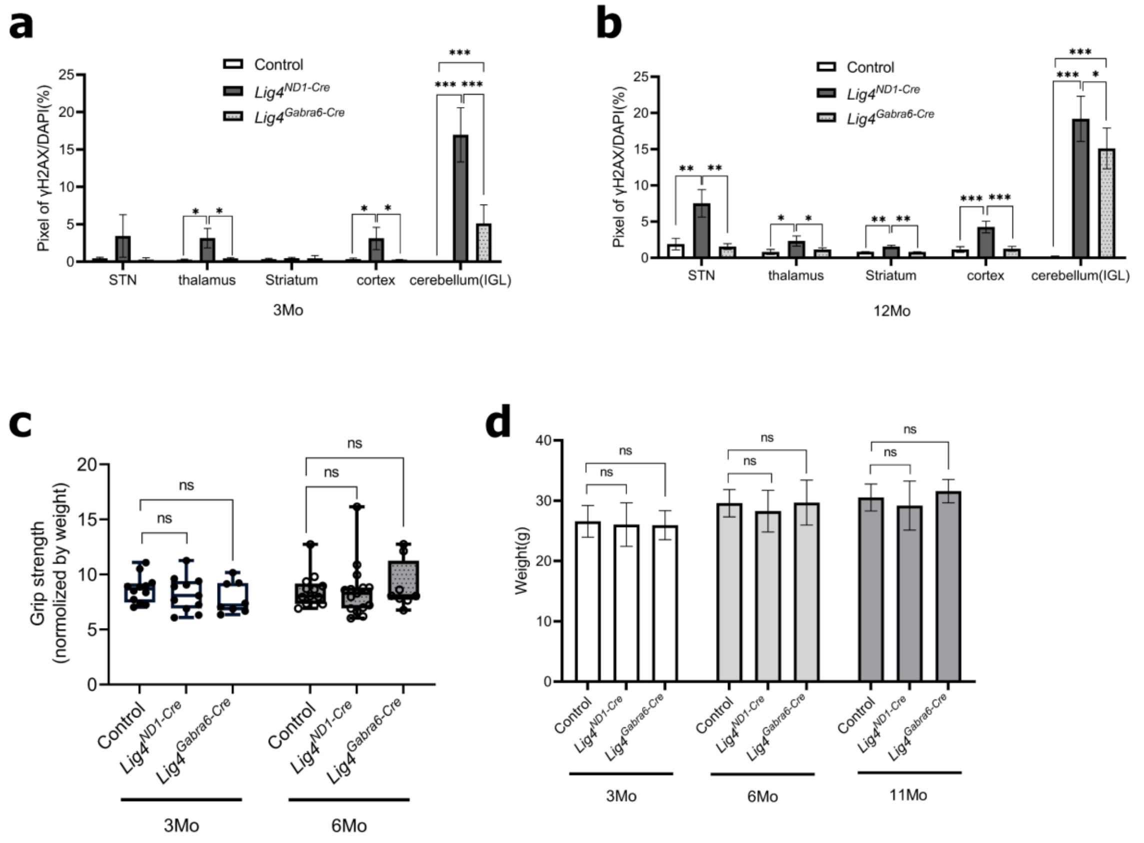


**Supplementary Fig. 7. Comparison of persistent DSBs and physical conditions of *Lig4* $^{\textit{ND1-Cre}}$ and *Lig4* $^{\textit{Gabra6-Cre}}$ mice.**

**a, b,** Quantification of γH2AX foci (relative pixel intensities of γH2AX and DAPI signals) in distinct brain regions of *Lig4* $^{ND1\text{-}Cre}$ and *Lig4* $^{Gabra6\text{-}Cre}$ mice at 6 mo (**a**) and 12 mo (**b**). Samples were collected from more than three independent experiments (n ≥ 3 mice per group). STN, subthalamic nucleus. **c,** Grip strength at 3 and 6 months of age (relative strength normalized to body weight; n > 10 for control and *Lig4* $^{ND1\text{-}Cre}$; n = 8 for *Lig4* $^{Gabra6\text{-}Cre}$). **d,** Body weight at three different ages (n ≥ 10 mice per group). Data represent mean ± s.d.; $*p < 0.05$, $**p < 0.01$, $***p < 0.001$; two-tailed unpaired t-tests.
